# A cell-nonautonomous heme acquisition pathway enables erythroid hemoglobinization under stress

**DOI:** 10.64898/2026.02.10.705195

**Authors:** Audrey Belot, Andrew Rock, Sohini Dutt, Gia Haemmerle, Amaury Maros, Xiaojing Yuan, Satoru Otsuru, David Bodine, Iqbal Hamza

**Affiliations:** Center for Blood Oxygen Transport and Hemostasis, Department of Pediatrics, School of Medicine, University of Maryland, Baltimore, Maryland, USA; Department of Animal and Avian Sciences, University of Maryland, College Park, Maryland, USA; Institute for Genome Sciences, University of Maryland School of Medicine, Baltimore, MD, USA; Department of Orthopedics, School of Medicine, University of Maryland, Baltimore, Maryland, USA; Genetics and Molecular Biology Branch, National Human Genome Research Institute, NIH, Bethesda, Maryland, USA

## Abstract

Heme, an iron-containing cofactor, is synthesized in mitochondria by an eight-enzyme pathway. Although cells were thought to manage heme autonomously, over 1,000 proteins contribute to its production, transport, and regulation. During terminal erythroid differentiation, mitochondria are discarded yet hemoglobin production continues, implying a cell-nonautonomous heme supply. We show that, under stress, erythroblasts import heme through the permease Heme Responsive Gene 1 (HRG1), which localizes to the plasma membrane and accumulates during stress erythropoiesis, the emergency program that expands red cell output. HRG1 loss impaired heme uptake, inhibited terminal erythroid differentiation, and caused anemia. In β-thalassemic mice, partial HRG1 loss reduces ineffective erythropoiesis, underscoring the importance of balanced heme import. These findings reveal intercellular heme sharing and identify HRG1 as a potential therapeutic target in hemoglobinopathies.

## Main Text

Over 90% of iron in the human body exists as heme, an iron-containing organic heterocyclic compound that is a vital cofactor responsible for diverse biological functions(*1*). In vertebrates, heme is synthesized by eight sequential enzymatic steps shared between the cytoplasm and mitochondria(*1*). A large number of effectors are essential for heme synthesis. These include intermediates from other metabolic pathways, such as succinyl-CoA, enzymatic cofactors, such as pyridoxal 5’-phosphate, and iron. Cells are thought to meet their heme requirements through their intrinsic ability to synthesize and regulate heme production, i.e. cell-autonomous regulation(*2*). However, a cell-nonautonomous heme communication system may exist in mammals – a concept largely unexplored. Converging evidence indicates vertebrates traffic heme between organs: cultured epithelial cells and macrophages release intact heme(*3, 4*); intravenous heme restores hepatic enzymes in porphyria(*5, 6*); zebrafish heme synthesis mutant embryos persist for 10-12 days presumably due to maternal heme(*7, 8*); monocytes acquire exogenous heme to form red-pulp macrophages(*9, 10*); and mammalian cells can incorporate and route exogenous heme to all major intracellular organelles (*11*). Given heme’s hydrophobicity and cytotoxicity, its movement across membranes must be tightly regulated through specific intra- and intercellular transport pathways(*12*). Still, the function and necessity of heme importers and exporters in mammals has been debated(*13*).

The majority of heme in the human body is within hemoglobin in red blood cells (RBCs)(*14-16*). During erythroid precursor development, heme synthesis and globin production are temporospatially coordinated to augment hemoglobin production. The iron importer transferrin receptor 1(*17*) and exporter ferroportin(*18*) are found in erythroid precursors to regulate iron levels for heme production. Much knowledge of heme metabolism in erythroblasts is restricted to heme synthesis and globin production, whereas the pathways responsible for heme transport and trafficking in erythroblasts are less well understood. Heme oxygenase 1 (HMOX1), which degrades heme to release iron, is expressed in erythroid precursors(*19*), raising questions as to why cells that are primed for heme production would degrade heme. Paradoxically, the maximal heme requirement occurs during late stages of erythroid maturation when the cells are systematically expelling their intracellular organelles, including the nucleus, Golgi, mitochondria, and secretory pathway, as they terminally differentiate(*20*).

For continued hemoglobin production, one plausible explanation is that erythroid cells satisfy their heme demands through a cell-nonautonomous mechanism. To explore this possibility, we performed single-cell RNA sequencing (scRNA-seq) and single-molecule fluorescence in situ hybridization (smFISH) on mouse erythroid precursors. These analyses revealed expression of the heme transporter Heme Responsive Gene 1 (*HRG1/SLC48A1*), a heme importer that is essential for heme-iron recycling by reticuloendothelial macrophages following erythrophagocytosis. Our findings reveal that HRG1 is essential for erythroid maturation under conditions of stress erythropoiesis, an emergency compensatory pathway that accelerates RBC production, largely in the spleen, when steady-state erythropoiesis in the bone marrow cannot meet physiological needs. HRG1 functions as a heme transporter and cycles between the plasma membrane and endocytic compartments. Loss of HRG1 in a β-thalassemia *Hbb*^*Th3/+*^ mouse model, characterized by heme overload within RBCs, improved ineffective stress erythropoiesis and anemia. Our genetic, cellular, and biochemical evidence supports a model in which erythroid precursors acquire heme through a cell-nonautonomous pathway mediated by HRG1.

## Results

### Erythroblasts express *HRG1*

We identified a role for *HRG1/SLC48A1* in heme transport in both *C. elegans* and mammals (*21, 22*). Mice genetically-depleted of HRG1 (*HRG1-KO*) are unable to respond to iron deficiency and have impaired erythroid maturation, conceivably due to *HRG1* deficiency and defective heme-iron recycling in reticuloendothelial macrophages(*23*). However, single-cell transcriptomics (scRNA-seq) revealed *HRG1* mRNA expression was highest within oligodendrocytes in the brain followed by megakaryocyte-erythroid progenitor (MEP) cells in the bone marrow (**Fig. 1A**) (*24*). Indeed, scRNA-seq showed *HRG1* mRNA was expressed in low amounts in the bone marrow within progenitors - multipotent progenitors (MPP) to early erythroid progenitors (EEP) - at steady-state and after erythropoietin (EPO) injection(*25*). Furthermore, *HRG1* expression increased as erythroid progenitors matured into late erythroid precursors (committed erythroid progenitors, CEP, to erythroid terminal differentiation, ETD), and this increase was further accentuated by erythropoietin injection (**Fig. 1B**). To validate these findings, Ter119-expressing cells from the bone marrow of a WT mouse were sorted by flow cytometry (**Fig. S1A**), Ter119 being an erythroid-lineage marker associated with committed erythroid precursors. Erythroblasts are classified as five distinct populations from the most immature population I (proerythroblasts), population II (basophilic erythroblasts), population III (polychromatic erythroblasts), population IV (orthochromatic erythroblasts and reticulocytes), to the most mature population V (RBCs)(*26*). scRNA-seq on these cells confirmed the erythroid lineage (**Fig. S1B**) (*26*). β-globin (*Hbb-bt*) mRNA was expressed throughout the entire UMAP (Uniform Manifold Approximation and Projection), indicating it is present in all erythroblast populations, unlike the expression of glycophorin A (*Gypa*, an erythroid marker) and transferrin receptor 1 (*Tfr1*), which were restricted to some populations (**Fig. 1C**). Most importantly, *HRG1* mRNA (*Slc48a1*) was present in all erythroblast populations (**Fig. 1C and D**). Proteomic data of human CD34-containing cells differentiated into the erythroid lineage(*27*), revealed detectable HRG1 protein in low amounts in progenitors 1 and 2, with higher amounts as the cells matured from proerythroblasts to orthochromatic erythroblasts (**Fig. 1E**). To spatially locate *HRG1* mRNA in the bone marrow, we used RNAscope, which enabled single-molecule visualization of RNA. *HRG1* was highly expressed in *SpiC-*containing cells, characteristic of central nurse macrophages, and in surrounding erythroblasts, typical of erythroblastic islands(*22, 28*) (**Fig. 1F**). These results indicate that HRG1 is expressed in erythroblasts.

**Figure 1.**
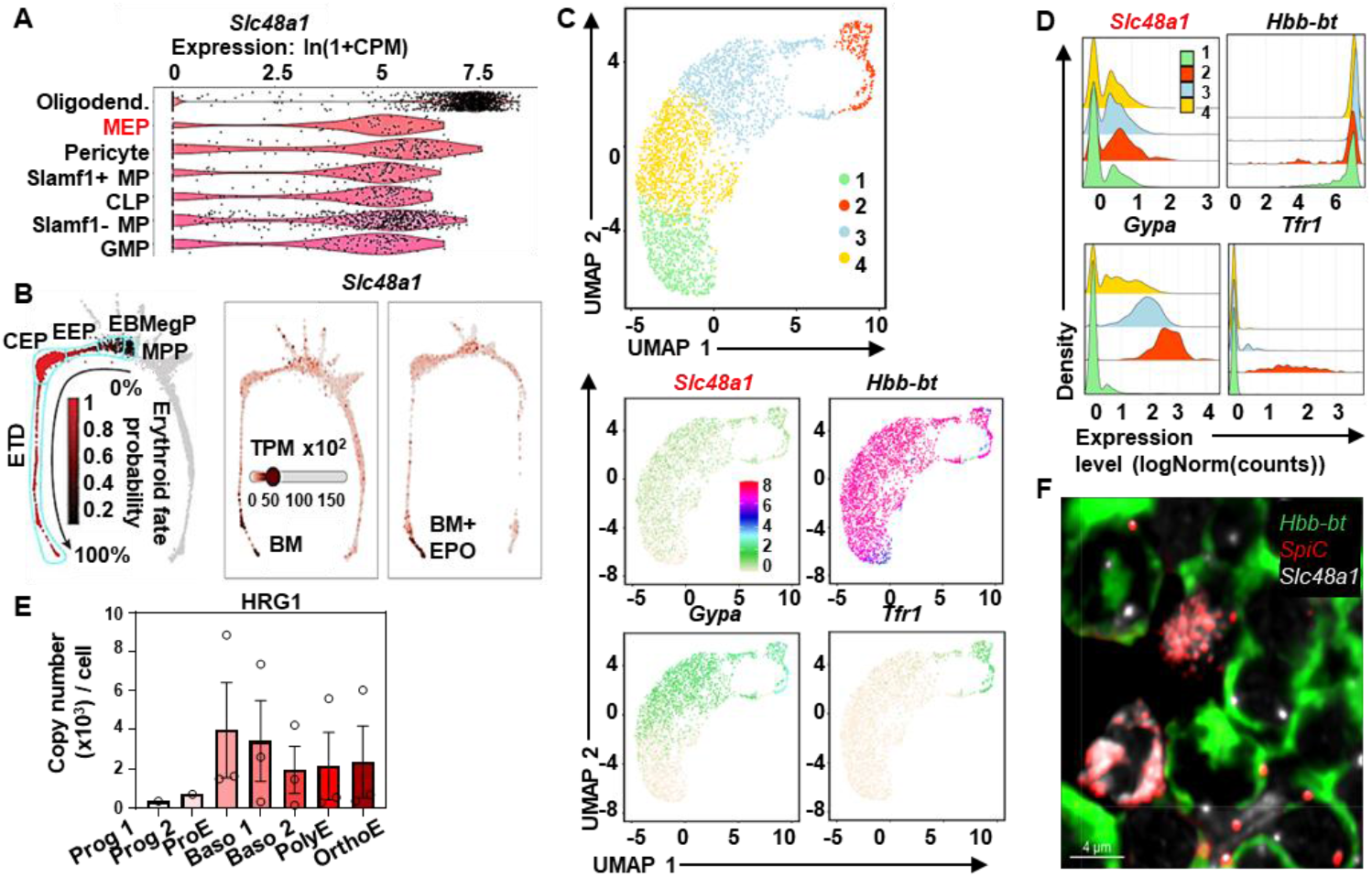
*Slc48a1*/HRG1 is present in erythroblasts at the mRNA and protein levels. **(A)** Violin plot extracted from Tabula Muris(19) of *Slc48a1* (HRG1) in all mouse tissues showing the top 7 cell types and tissues expressing *Slc48a1*/HRG1. Oligodend.: Oligodendrocyte (Brain), MEP: Megakaryocyte Erythroid progenitors (bone marrow), Pericyte: Pericyte (Brain), Slamf1+ MP: Slamf1 positive multipotent progenitor cell (bone marrow), CLP: Common Lymphoid Progenitor, Slamf1-MP: Slamf1 negative multipotent progenitor cell (bone marrow), GMP: Granulocyte Monocyte Progenitor cell (bone marrow). **(B)** Data extracted from Tusi et al(20) and visualized for *Slc48a1*/HRG1 in bone marrow (BM) cells in control or EPO (erythropoietin) conditions. MPP: multipotent progenitors; EBMegP: erythroid–basophil–megakaryocyte-biased progenitors; EEP: early erythroid progenitors; CEP: committed erythroid progenitors; ETD: erythroid terminal differentiation. **(C)** Single-cell RNA-seq of Ter119-containing cells from the bone marrow of a WT mouse. UMAP shows cell grouping from 1 to 4, as well as gene expression of *Slc48a1* (HRG1), *Hbb-bt* (β-globin), *Gypa* (glycophorin A), and *Tfr1* (Transferrin receptor 1) in the Ter119 population. **(D)** Ridge plot of *Slc48a1, Hbb-bt, Gypa*, and *Tfr1* gene expression in the WT Ter119 population from the single-cell RNA-seq. **(E)** Proteomic data extracted from Gautier et al(22) from CD34-containing cells differentiated cells in erythroblasts showing HRG1 expression across different stages of maturation. **(F)** mRNA in situ hybridization of WT bone marrow phlebotomized twice with mRNA probes for *Hbb-bt* (green), *SpiC* (red) and *Slc48a1*/HRG1 (green) and Tfr1 antibodies. Scale bars as indicated.

### *HRG1* deficiency induced severe anemia during stress erythropoiesis

To determine if *HRG1* has a role in erythropoiesis, we treated *HRG1-KO* mice with phlebotomies (blood withdraw) or erythropoietin to induce stress erythropoiesis (**Fig. S2A**). Under steady-state, *HRG1-KO* mice have a similar spleen size to WT, but spleens were larger in phlebotomized and erythropoietin-injected mice, suggesting ineffective stress erythropoiesis in *HRG1-KO* mice (**Fig. 2A**); liver and kidney sizes were unaffected (**Fig. S2B and C**). While phlebotomies caused hematocrit and RBC count to decrease significantly in WT mice, the changes were greater in the *HRG1-KO* mice with lower RBC count and hematocrit (**Fig. 2B and C**; **Table S1**).

**Figure 2.**
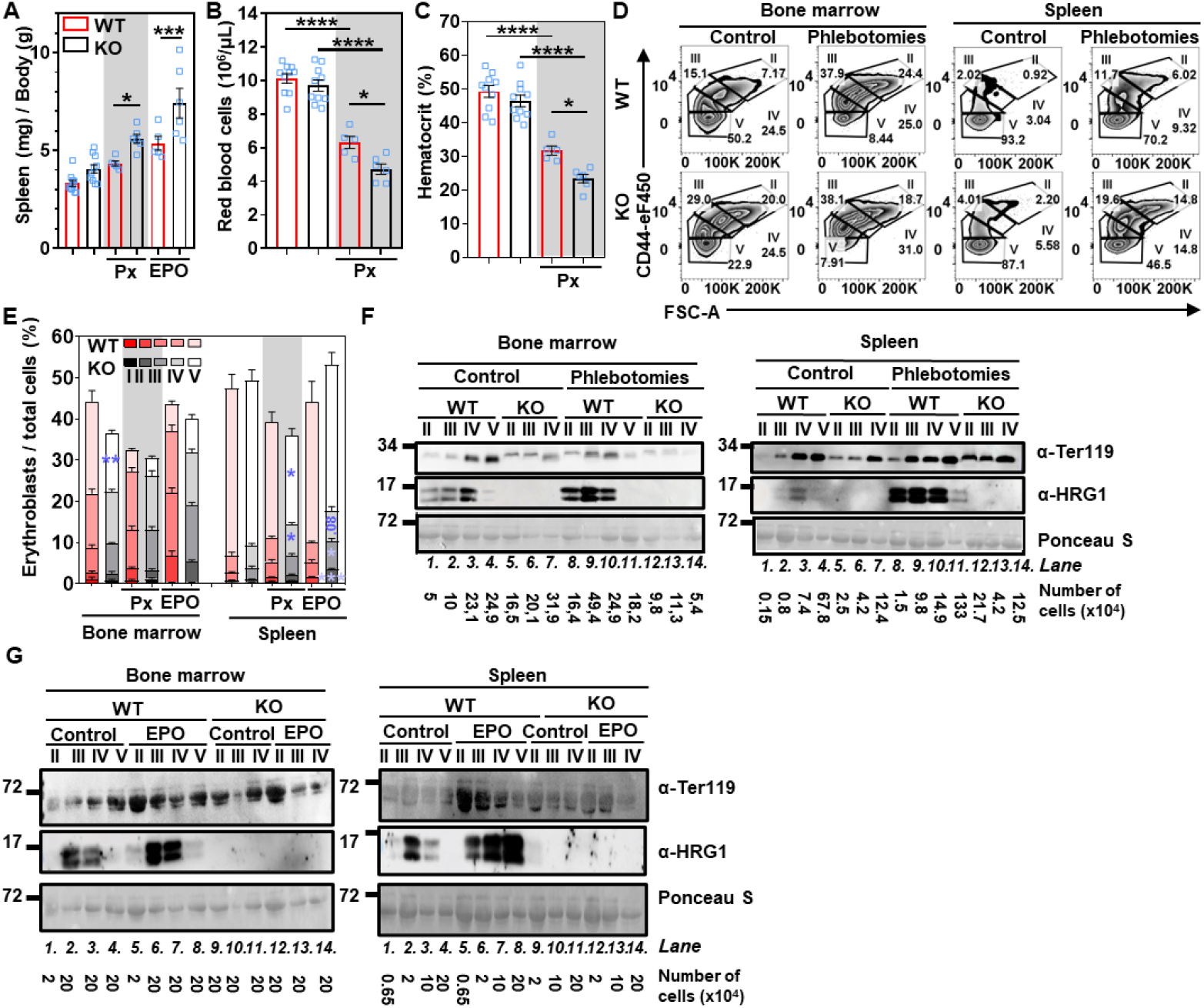
HRG1 is induced by stress erythropoiesis, and its deletion leads to more severe anemia during high requirements of heme. **(A)** The ratio of spleen weight normalized to the body weight of WT and *HRG1-KO* male mice (open blue square) in control, phlebotomies (Px), and erythropoietin (EPO) conditions. Error bars represent mean ± SEM, n=5-12 mice/group. **p<0.01, ***p<0.001 (One-way ANOVA, Sidak’s multiple comparison test). Blood parameters from WT and *HRG1-KO* male mice (open blue square), **(B)** red blood cell count, and **(C)** hematocrit with or without phlebotomies (Px). Error bars represent mean ± SEM, n=5-10 mice/group. *<0.05, ***p<0.001, ****p<0.0001 (One-way ANOVA, Sidak’s multiple comparison test). **(D)** Representative flow cytometry plots of the different stages of erythroblasts in WT and *HRG1-KO* mice in control or phlebotomized conditions. Cells were gated for CD44 and cell size (FSC-A, forward scatter area). **(E)** Flow cytometry analysis of the proportion of erythroid precursors in the live single cell population in the bone marrow and the spleen from WT and *HRG1-KO* male mice in control, phlebotomies (Px), and EPO conditions. WT and HRG1-KO mice from the same condition for each erythroid population were compared using a t-test. *p<0.05, **p<0.01, ***p<0.001.Immunoblots of Ter119 and HRG1 from populations II to V from **(F)** phlebotomized or **(G)** injected with EPO of WT and *HRG1*-KO male mice from the bone marrow and the spleen. The number of cells loaded per well is indicated. These immunoblots are representative of biological replicates, which were at least reproduced twice.

To elucidate the mechanisms underlying how *HRG1* deficiency causes fewer circulating RBCs after phlebotomies, erythroid populations in the bone marrow and spleen of male mice were quantified by flow cytometry. Population I expresses Ter119 at a medium abundance, while its abundance is high in populations II to V. Although steady-state blood parameters (RBC count and hematocrit) were comparable between WT and *HRG1-KO* adult mice (**Fig. 2B and C**), flow cytometry revealed a significant reduction in the most mature erythroid population (population V) among total viable bone marrow cells (**Fig. 2D, left panel; Fig. 2E**). This decrease was attributable to increased apoptosis of Ter119-positive erythroblasts (**Fig. S2D and E**). After phlebotomies or erythropoietin treatment, the inhibition of bone marrow erythroid differentiation was exacerbated in *HRG1-KO* mice with a concomitant impairment in expansion of population IV (orthochromatic erythroblasts and reticulocytes) and apoptosis of Ter119-containing cells in the spleen (**Fig. 2D right panel, 2E; Fig. S2D and E**). In female mice, no significant difference was observed, and there were greater variations in splenomegaly. There was no change in liver and kidney weights and induction of erythroid populations II to IV in the bone marrow and the spleen (**Fig. S2F-J**).

This observation aligns with previous findings, such as the regulation of erythropoiesis by sexual hormones, particularly androgens (*29*), which indicates a differential response in males and females.

At steady-state, HRG1 protein was detected in populations II to IV in the bone marrow (**Fig. 2F, 2G, lanes 1-3**) and populations III and IV in the spleen (**Fig. 2F and G, lanes 2-3; Figure S2K and L**). In contrast, Ter119 was detectable in populations II to V in the bone marrow and spleen. During stress erythropoiesis, HRG1 was induced in populations II to IV in both the bone marrow (**Fig. 2F, lanes 8-10**) and spleen (**Fig. 2G, lanes 5-7**). Altogether, these findings indicate that the absence of HRG1 impaired erythropoiesis, leading to increased apoptosis of erythroid precursors and, ultimately, anemia.

### HRG1 is important for terminal erythroid differentiation and imports exogenous heme to support erythroid maturation

To investigate the role of HRG1 in erythroid precursors, we used K562 cells, a human erythroleukemia cell line. These cells can be differentiated by treatment with sodium butyrate (SB) into the erythroid lineage until the orthochromatic stage(*30-32*). We generated several *HRG1-KO* cell lines using CRISPR/Cas9 by either deleting exon 1 (KO) or the sequence between exons 1 and 3 (KO#2 and KO#3), and each KO clone was confirmed by sequencing to identify the precise genetic lesion (**Fig. S3A**). K562 cells treated with sodium butyrate showed increased HRG1 protein (**Fig. 3A**), accumulated more heme (**Fig. 3B**), and produced more benzidine-positive hemoglobinized cells (**Fig. 3C**). By contrast, *HRG1-KO* cells failed to accumulate heme (**Fig. 3B**) or fully make hemoglobin regardless of varying the concentrations of either sodium butyrate (**Fig. 3C, Fig. S3B**) or hemin, another differentiation factor (**Fig. 3D, Fig. S3C**). Immunofluorescence microscopy identified endogenous (**Fig. 3E**) or transfected HRG1 (**Fig. S3D**) on vesicular and plasma membranes, partially colocalizing with transferrin receptor 1 (Pearson correlation coefficient = 0.4579)(*33*).

**Figure 3.**
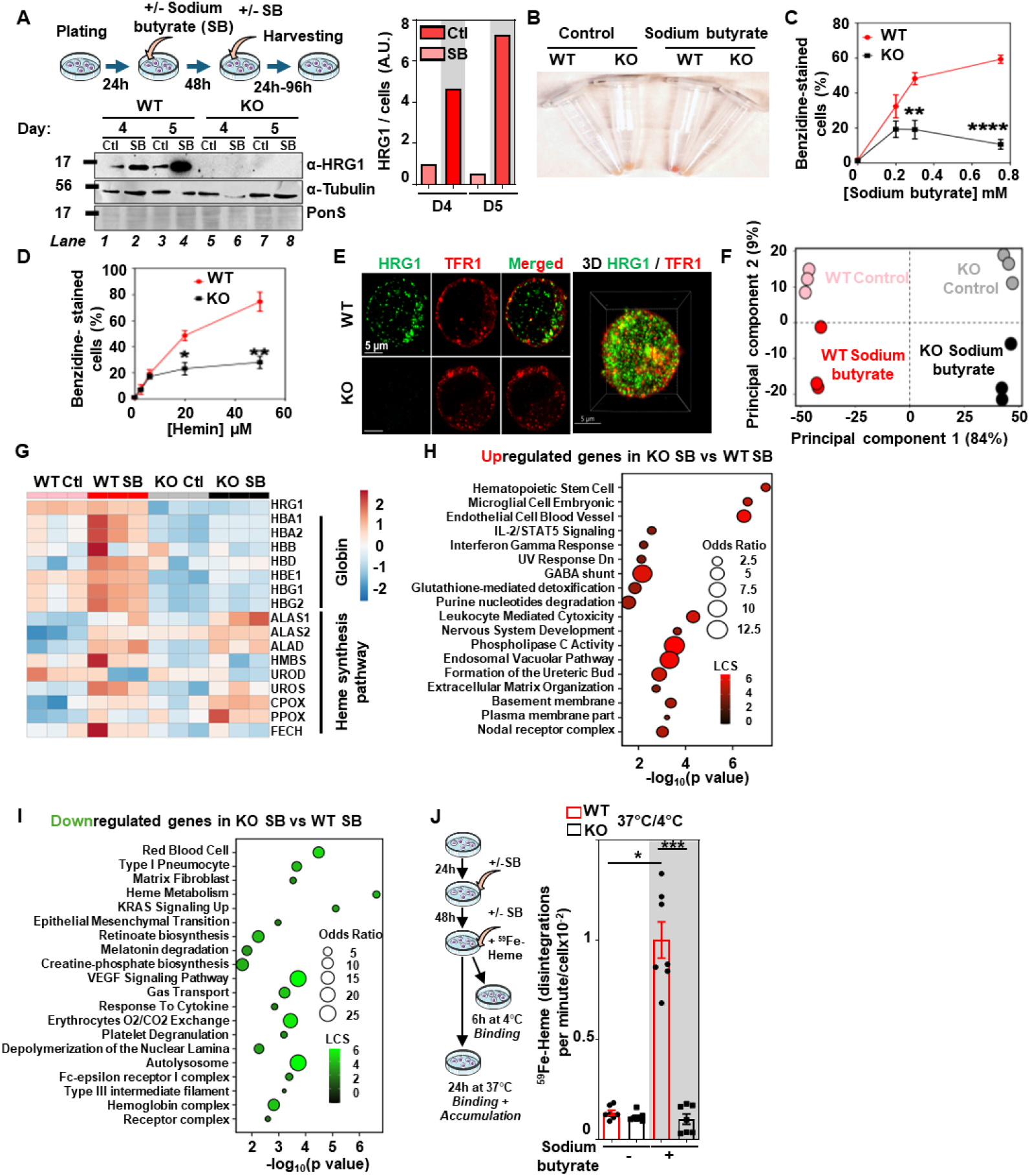
HRG1 imports exogenous heme for erythroblasts to fully hemoglobinize. **(A)** The schematic represents an experimental design. Briefly, K562 WT and *HRG1*-KO K562 cells were plated, treated with 0.75 mM sodium butyrate (SB), treated again 48h later, and finally harvested at different time points (between 24 hours and 96 hours) after the second treatment. Immunoblot of HRG1 and α-tubulin from WT and *HRG1*-KO K562 cells 96 hours after the second treatment of SB. Blots were quantified (right panel). The experiment was independently replicated biologically twice. **(B)** Cell pellets of WT and *HRG1*-KO K562 cells, after being treated twice with 0.75 mM SB and harvested 96 hours after the second treatment. WT (red circle) and *HRG1*-KO (black square) K562 cells were treated twice with different concentrations of **(C)** sodium butyrate or **(D)** hemin, harvested 96 hours after the second treatment, and stained with o-dianisidine (benzidine). Error bars represent mean ± SEM from 3 biological replicates comprising at least 2 technical replicates. WT and HRG1-KO cells were compared at each time point using a t-test *p<0.05, **p<0.01, ***p<0.001, ****p<0.0001. (**E**) Immunofluorescence of HRG1 (green) and TFR1 (transferrin receptor 1, red) WT and *HRG1*-KO K562 cells treated twice with 0.75 mM sodium butyrate. Pearson coefficient between HRG1 and TFR1 of 0.4579. Scale bars as indicated. (**F**) Principal components analysis (PCA) of WT and *HRG1*-KO K562 cells treated or not with 0.75 mM SB on three biological replicates for each condition. (**G**) Heatmap of heme biosynthesis pathway and globin genes in WT and *HRG1*-KO K562 cells in control or SB-treated conditions. Enrichment analysis plots of (**H**) the 1232 genes that are upregulated or (**I**) 998 genes that are downregulated in *HRG1*-KO K562 cells SB compared to WT SB. (**J**) The schematic represents the experimental design. WT (red circle) and *HRG1*-KO (black square) K562 cells were plated, treated with 0.75 mM SB 24 hours later, treated again 48 hours later with SB and with ^59^Fe-Heme, and incubated either at 4°C for 6 hours or 37°C for 24 hours. Error bars represent mean ± SEM from 3 biological replicates, comprising 2-3 technical replicates.*p<0.05, ***p<0.001 (One-way ANOVA, Sidak’s multiple comparison test).

To confirm that HRG1 loss impaired both hemoglobin accumulation and red cell maturation, we performed differential gene expression analysis using RNA-seq data from control and differentiated WT and *HRG1-KO* K562 cells. Principal component analysis (PCA) clustered the four conditions based on the genotype (84% of the variance) and the sodium butyrate treatment (9% of the variance; **Fig. 3F**). Out of 391 genes dysregulated, 374 genes were differentially regulated between WT and *HRG1-KO* in the presence of sodium butyrate (**Fig. S3E**). Treatment with sodium butyrate did not change the abundance of *HRG1* mRNA **(Fig. 3G)** even though protein amounts increased **(Fig. 3A)**, indicating that *HRG1* must be regulated post-transcriptionally. Gene expression heatmaps revealed that transcription of globin genes was significantly upregulated in WT sodium butyrate compared to WT control (*HBA1* and *-2, HBD, HBE1, HBG1* and *-2*), but strongly downregulated in *HRG1-KO* and not induced by sodium butyrate (**Fig. 3G**). Heme biosynthesis pathway genes showed increased expression in cells treated with sodium butyrate in both WT and *HRG1-KO* cells (*ALAS1, ALAS2, ALAD, UROS, CPOX, PPOX*), with the exception of *HMBS, UROD*, and *FECH*. Thus, cells lacking *HRG1* appear to be unable to upregulate *FECH*, the final enzyme in the heme biosynthesis pathway, in response to hemoglobinization, further exacerbating heme deficiency.

As heme and iron metabolism are strongly intertwined, we analyzed iron metabolism pathway genes (**Table S2**). Several genes implicated in iron transport (*TFR2*), storage (*FTL*), and ferroptosis (*SLC38A1*) were downregulated in *HRG1-KO*-treated cells. As expected, gene set enrichment analysis (GSEA) showed significant activation of genes involved in the erythroid lineage, heme metabolism, heme biosynthesis, oxygen transport, and the hemoglobin complex in WT cells treated with sodium butyrate (**Fig. S3F**). By contrast, sodium butyrate upregulated hematopoietic stem cell genes to a greater extent in *HRG1-KO* cells than it did in WT (**Fig. 3H**) and downregulated RBCs development, heme metabolism, and the hemoglobin complex genes **(Fig. 3I)**. These findings suggest that *HRG1* deficiency causes failure in terminal erythroid differentiation.

To directly measure heme transport, we used radioactive [^59^Fe]heme tracer. K562 cells were incubated with [^59^Fe]heme for 24 hours after treatment with sodium butyrate. WT cells treated with sodium butyrate showed >10-fold increase in [^59^Fe]heme accumulation that was attenuated in KO cells (**Fig. 3J, Fig. S3G and H**). In comparison, WT cells showed ≈3-fold more ^59^Fe accumulation and binding than did KO cells (**Fig. S3I-S3K**). Thus, HRG1 is localized to the plasma membrane of erythroid precursors, mediates heme import into the cells, and is essential for efficient hemoglobin production.

### *HRG1-GFP* mouse model reveals stage-specific regulation of HRG1 during stress and iron-deficiency anemia

To modulate and quantify HRG1 abundance, we generated an *HRG1-GFP* mouse model in which the green fluorescent protein (GFP) was inserted into the *HRG1* locus through homology-directed repair using CRISPR/Cas9 (**Fig. 4A**), resulting in the expression of an HRG1-GFP chimeric protein. The *HRG1-GFP* mouse allows the direct quantification of HRG1 temporospatial expression and localization in erythroid populations by flow cytometry (**Fig. S4A)**. FlowSight, a flow cytometer combined with a microscope, showed HRG1 localized with Ter119 and CD44, which mark the plasma membrane and vesicular compartments, respectively, in populations II to IV of the bone marrow, confirming our colocalization studies in K562 cells (**Fig. 4B, S4B**). At steady-state, HRG1-GFP was detected in populations II to IV in the bone marrow, and undetectable in populations II and III in the spleen due to the absence or very low abundance of these populations (**Fig. 4C** versus **2F**), corroborating that erythropoiesis in *HRG1*^*GFP/+*^ mice was normal. Upon erythropoietin-injection or phlebotomy, the number of HRG1-containing cells as well as the amount of HRG1 per cell were significantly greater in populations II to IV in both the bone marrow and spleen (**Fig. 4C and D, S4C**).

**Figure 4.**
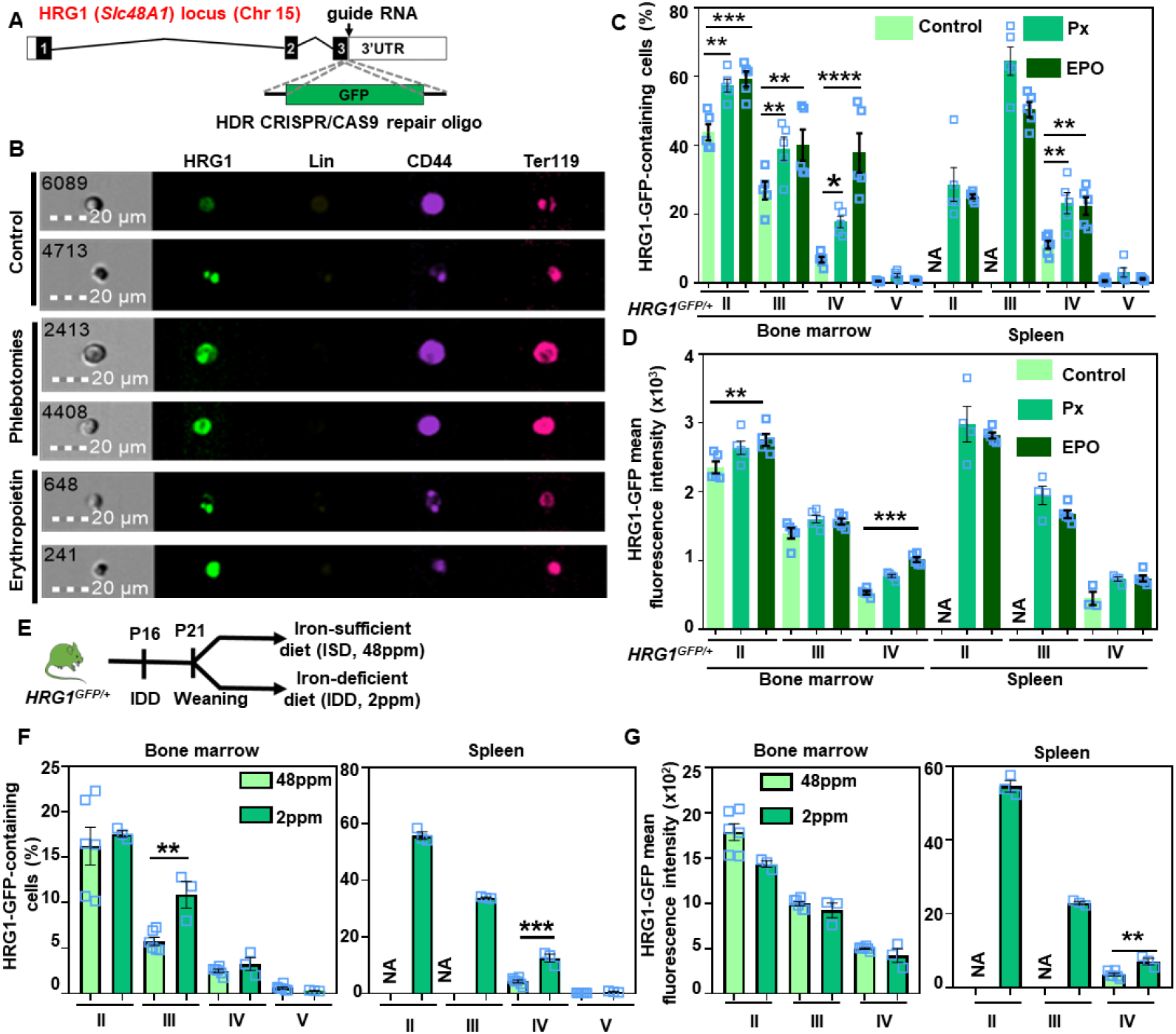
HRG1 is induced during iron deficiency anemia. **(A)** Schematic of HRG1-GFP translational fusion mouse, which was created by recombining turboGFP into exon 3 of the HRG1 locus. (**B**) Representative microscopy images of bone marrow erythroblasts of HRG1^GFP/GFP^ mice in control, phlebotomy, and erythropoietin conditions using FlowSight. (**C**) Percentage of HRG1-GFP-containing cells and (**D**) HRG1-GFP mean fluorescence intensity in each erythroid precursor population in *HRG1*^*GFP/+*^ control, phlebotomized (Px), or EPO-injected male mice (open blue square), as previously described in Figure S2A. Error bars represent mean ± SEM from n=5 mice/group. **p<0.01, ***p<0.001, ****p<0.0001 (each population from control and treated conditions from either bone marrow or spleen was compared by t-test). (**E**) HRG1^GFP/+^ male mice were fed an iron-deficient diet (IDD, 2ppm of iron) at P16, weaned at P21 on either an iron-sufficient diet (ISD, 48ppm of iron) or IDD, and the experiment was performed at 8 weeks old. (**F**) Percentage of HRG1-GFP-containing cells and (**G**) HRG1-GFP mean fluorescence intensity (MFI) in each erythroid precursor population in HRG1^GFP/+^ male mice (open blue square) fed an iron-deficient diet or an iron-sufficient diet. Error bars represent mean ± SEM from n=3-5 mice/group. **p<0.01, ***p<0.001 (each population from IDD and ISD from either bone marrow or spleen was compared by t-test).

To investigate whether HRG1 is differentially regulated, *HRG1*^*GFP/+*^ mice were fed an iron-deficient diet (IDD, 2ppm of iron) starting at P16, and at P21 weaned on either an iron-sufficient (ISD, 48ppm of iron) or IDD prior to sacrifice at 8 weeks (**Fig. 4E**). This treatment causes anemia as a result of iron deficiency. IDD induced a drop in RBC count and hematocrit (**Fig. S4D and E**) and increased stress erythropoiesis (**Fig. S4F**). Although HRG1-positive cells were significantly more abundant in population III within the bone marrow, both the number of HRG1-expressing cells and the amount of HRG1 expression per cell were markedly higher in populations II to IV in the spleen (**Fig. 4F and G**). These findings further support a role for HRG1 in erythroid maturation and in stress erythropoiesis.

### HRG1 as a genetic modifier of β-thalassemia

To investigate HRG1 functions in hemoglobinopathies, we crossed the β-thalassemia mice [*Hbb*^*th3/+*^ (Th3/+)] with either *HRG1-GFP* (G/+) to obtain Th3/+;G/+ or *HRG1-KO* (KO) to obtain either Th3/+;HT (heterozygous) or Th3/+;KO (null) (**Fig. S5A**). β-thalassemia is caused by mutations and deficiencies in the β-globin gene, resulting in poor growth, hemolytic anemia, and ineffective stress erythropoiesis. Because the defects in β-globin chains result in the accumulation of heme, α-globin, hemichromes, and reactive oxygen species (ROS)(*34*), we reasoned that heme import by HRG1 could be a genetic modifier in β-thalassemia.

*Hbb*^*th3/+*^mice had over 2-fold more HRG1-GFP-containing cells in populations II to IV in the bone marrow (**Fig. 5A**). The spleen showed a 20-fold increase in cells expressing HRG1-GFP. The amount of HRG1-GFP per cell increased significantly in populations IV and V in the bone marrow, and populations II to IV in the spleen of Th3/+;G/+ mice (**Fig. 5B**). These findings confirm that HRG1 accumulates in erythroblasts during ineffective stress erythropoiesis.

**Figure 5.**
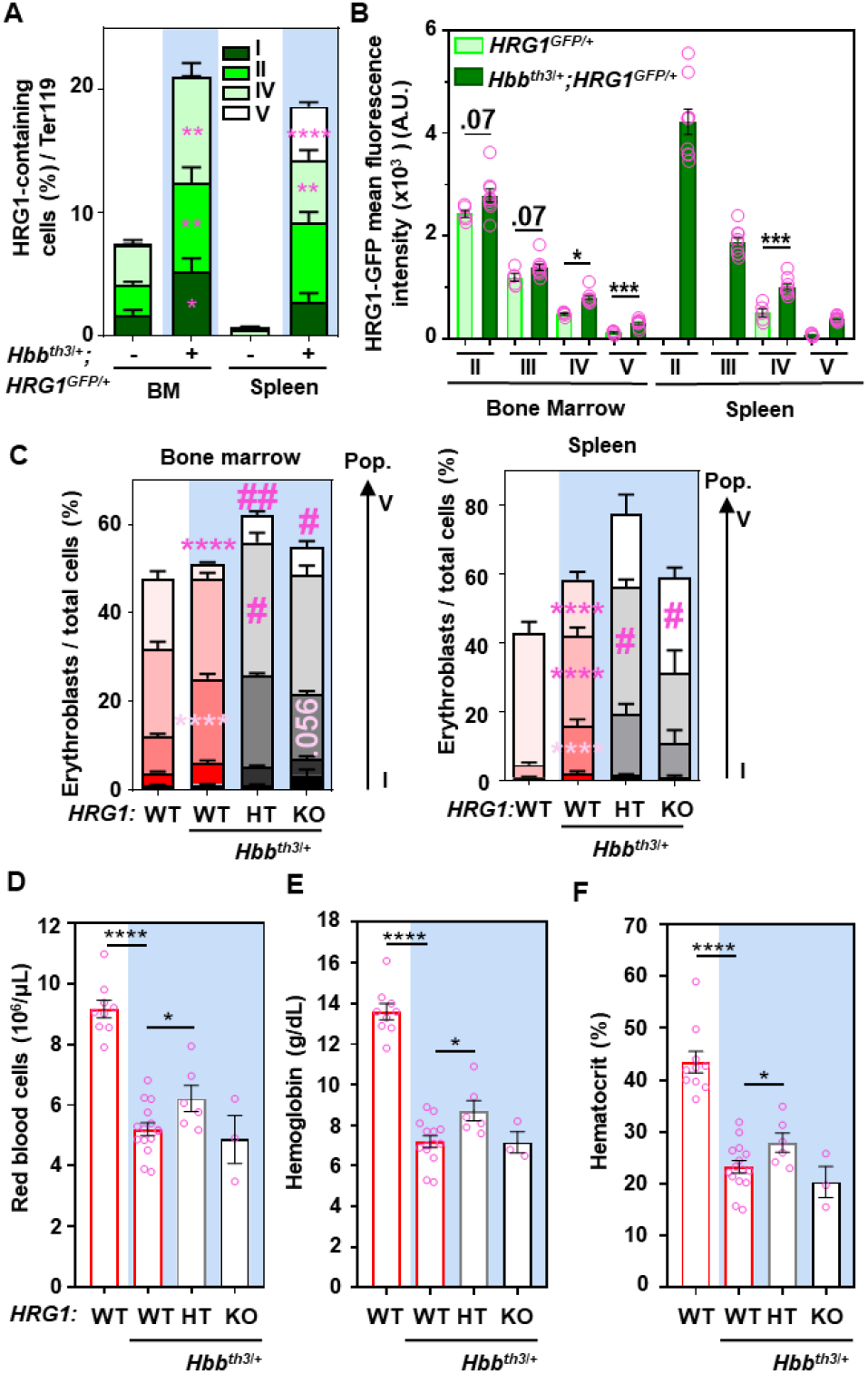
HRG1 is highly induced in erythroblasts in a β-thalassemic mouse model. **(A)** Percentage of HRG1-GFP-containing cells and **(B)** HRG1-GFP mean fluorescence intensity (MFI) in populations II to IV in Ter119-positive cells from the bone marrow (BM) and the spleen in *HRG1*^*GFP/+*^ (-) and *Hbb*^*Th3/+*^*;HRG1*^*GFP/+*^ (+) female mice (open pink circles). Error bars represent mean ± SEM, n=5-9 mice/group. *p<0.05, **p<0.01, ***p<0.001, ****p<0.0001 (each population from WT and *Hbb*^*th3/+*^ from either bone marrow or spleen was compared by t-test. **(C)** Flow cytometry analysis of the proportion of erythroid precursors in the single cell population in the bone marrow (left) and the spleen (right). Error bars represent mean ± SEM, n=4-12 mice/group. Each population within the same condition was compared using a t-test between WT and *Hbb*^*th3/+*^*;WT* mice. *p<0.05, **p<0.01, ***p<0.001. #<0.05, ##p<0.01: each population from *Hbb*^*th3/+*^and *Hbb*^*th3/+*^*;HT or KO* from either bone marrow or spleen was compared by t-test. Blood parameters from WT, *Hbb*^*th3/+*^, *Hbb*^*th3/+*^*;HRG1-HT, and Hbb*^*th3/+*^*;HRG1-KO* female mice (open pink circles) **(D)** red blood cells, **(E)** hemoglobin, and **(F)** hematocrit. Error bars represent mean ± SEM, n=3-10 mice/group. *<0.05, ***p<0.001, ****p<0.0001 (t-test between *Hbb*^*th3/+*^ and other groups). HT: HRG1 heterozygote.

To elucidate the role of *HRG1* in *Hbb*^*th/3+*^ erythroid precursors, we used the Th3/+;KO. We reasoned that deleting *HRG1* might improve ineffective stress erythropoiesis in *Hbb*^*th/3+*^ as heme import by HRG1 might cause further cytotoxicity. *Hbb*^*th3/+*^mice showed splenomegaly (**Fig. S5B**) and ineffective stress erythropoiesis in the bone marrow and spleen with an increase in populations II to IV and a decrease in population V (**Figure 5C**) with a concomitant anemia (**Fig. 5D-5F)**. However, deleting one allele of *HRG1* (Th3/+;HT) increased erythropoiesis significantly in both the bone marrow and the spleen (**Fig. 5C**), with a significant increase in populations IV and V in the bone marrow and population IV in the spleen. This was accompanied by an improvement in anemia with a significant increase in RBC count, hemoglobin, and hematocrit (**Fig. 5D-5F, Table S3)**.

Even though the percentage of erythroblasts increased by a third in the spleen (**Fig. 5C**, WT vs Th3/+;HT), splenomegaly was unchanged, as well as liver and kidney weights (**Fig. S5B-S5D**). By contrast, Th3/+;KO mice showed a significant increase in population V in the bone marrow and the spleen (**Fig. 5C**) but no improvement in anemia (**Fig. 5D-5F**). These findings indicate that complete loss of *HRG1* in both, recycling macrophages and erythroblasts, leads to pleiotropic effects, potentially obscuring any therapeutic benefit in β-thalassemia. Thus, fine-tuning HRG1 may be advantageous by mitigating stress erythropoiesis and improving anemia.

## Discussion

Intracellular heme is thought to be dictated by mitochondrial heme synthesis and degradation. However, as erythroid precursors mature, they systematically lose their organelles, including the mitochondria, even though hemoglobinization continues unabated(*35*). How can erythroid precursors maximally hemoglobinize when they lack functional mitochondria for heme synthesis? Comparing heme and iron metabolism pathways, the iron importer transferrin receptor-1(*17*) and exporter ferroportin(*18*) are both found in erythroid precursors to coordinate iron transport with heme synthesis. Although counterintuitive, iron export during terminal erythroid differentiation becomes necessary when cells have excess iron but lose the capability to synthesize heme(*18*). The existence and role of heme importers and exporters in mammals have been controversial(*13*). While, FLVCR1 was identified as a heme exporter in early erythroid precursors(*36*), the identity of a heme importer in these cells was missing. We demonstrate that the transmembrane heme permease HRG1/SLC48A1 imports heme in erythroid precursors and accumulates during stress erythropoiesis (**Fig. S5E**). Loss of HRG1 results in anemia and stress erythropoiesis with a failure in terminal erythroid differentiation.

Our findings demonstrate that HRG1 is essential for erythropoiesis under conditions of high heme demand, such as anemia and stress erythropoiesis. HRG1 is expressed in murine erythroblasts and human erythroleukemia cells, and exhibits a vesicular and plasma membrane localization, which positions the transporter for import of exogenous heme. This is supported by the presence of tyrosine- and di-leucine-based sorting signals on the C-terminus of HRG1(*21, 37*), motifs known to be responsible for vesicular and plasma membrane localization. HRG1 accumulates during stress erythropoiesis in basophilic erythroblasts in the bone marrow and spleen, and its deletion leads to apoptosis of erythroid precursors due to insufficient hemoglobinization resulting in anemia.

The apparent import of exogenous heme by HRG1 raises the question of its origin. Heme could come from nurse macrophages within the erythroblastic islands through direct cell-cell contact involving a macrophage heme exporter such as FLVCR1. However, studies have also implicated FLVCR1 in choline import(*38-40*). Another plausible pathway could be through macrophage-derived extracellular vesicles, which may carry macromolecules and small metabolites(*41-44*).

We show that HRG1 could be a genetic modifier of β-thalassemia, a disease characterized by heme-induced cytotoxicity in the erythroid compartment. Indeed, HRG1 is highly induced in erythroblasts in a β-thalassemia mouse model, but even though fully ablating HRG1 did not improve the anemia, deleting one allele of *HRG1* improved ineffective stress erythropoiesis in the bone marrow and spleen, and consequently, anemia. Fine-tuning HRG1 expression might be beneficial in anemia-related diseases, such as β-thalassemia and porphyrias.

## Supporting information

All Supplementary Files

## Acknowledgments

We thank Lisa Garrett and the NHGRI genomics core for their assistance in generating the HRG1-GFP mice, Fernanda Laranjeira Da Silva for the initial genotyping of the HRG1-GFP mice, and Jianbing Zhang for generating the HRG1-KO K562 cell line; Harry Dailey for critical discussions and reading of the manuscript; Shilpa Kumar and the Confocal Microscopy Core at the Center for Innovative Biomedical Resources, University of Maryland School of Medicine Baltimore; the Flow Cytometry Shared Service of the University of Maryland Marlene and Stewart Greenebaum Comprehensive Cancer Center for the use of the LSR2, Aurora Cytek3, BD Aria II. Flowsight, Maryland Genomics at the Institute for Genome Sciences, University of Maryland School of Medicine, for performing the single-cell RNA sequencing. The [^59^Fe] isotope used in this research was supplied by the U.S. Department of Energy Isotope Program, managed by the Office of Isotope R&D and Production. Illustrations adapted from Servier Medical Art (https://smart.servier.com) used under CC BY 4.0 license.

This work was supported by funding:

National Institutes of Health grant DK125740 (IH)

National Institutes of Health grant DK134583 (IH)

Co-Operative Center for Excellence in Hematology type B grant #10055841-22-UM (AB)

Cooley’s Anemia Foundation grant #3006495 (AB)

The Maryland Department of Health’s Cigarette Restitution Fund Program – CH-649-CRF (Flow cytometry shared service)

The National Cancer Institute - Cancer Center Support Grant (CCSG) - P30CA134274 (Flow cytometry shared service)

## Author contributions

Conceptualization: AB, DB, IH

Methodology: AB, AR, GH, SD, XY, AM, SO

Investigation: AB, IH

Visualization: AB, AM, XY

Funding acquisition: AB, IH

Supervision: IH

Writing – original draft: AB, IH

Writing – review & editing: All the authors

## Data and materials availability

All raw sequencing data generated in this study have been deposited in the NCBI Sequence Read Archive (SRA) under the BioProject accession PRJNA1366593. The corresponding BioSample and SRA run accessions for each sample are provided in Supplementary Table 4. Processed data have been deposited in the Gene Expression Omnibus (GEO) under the following accessions: GSE311281, containing the barcode-feature matrix for the single-cell RNA-seq sample, and GSE311284, containing the counts table for the bulk RNA-seq data from K562 cells. The code used for all analyses is available on GitHub and has been archived on Zenodo under the DOI https://doi.org/10.5281/zenodo.17676483.

