## Supplementary material for "A cell-nonautonomous heme acquisition pathway enables erythroid hemoglobinization under stress": All Supplementary Files

**The PDF file includes:**

Materials and Methods
Figs. S1 to S5
Tables S1 to S4
References (45-64)

#### Materials and Methods

##### Materials

| Antibodies and dyes | Source | Catalog number |
|---|---|---|
| Mouse monoclonal anti-tubulin | DSHB | #AA4.3 |
| Rat monoclonal anti-Ter119 | ThermoFisher | #14-5921-85 |
| Rabbit polyclonal anti-HRG1 | Pek et al.(*1*) | N/A |
| Goat anti-Rabbit IgG (H+L) Secondary Antibody, HRP | Invitrogen | #31460 |
| Goat anti-Rat IgG (H+L) Secondary Antibody, HRP | ThermoFisher | # 31470 |
| Ter119-APCeFluor780 | Invitrogen | #47-5921-82 |
| CD44-PECy7 | Biolegend | #103030 |
| CD44-eFluor450 | Invitrogen | #40-0441-82 |
| Gr-1-PE | Invitrogen | #12-5931-83 |
| CD4-PE | Invitrogen | #12-0041-82 |
| CD8a-PE | Invitrogen | #12-0081-83 |
| B220-PE | Invitrogen | #12-0452-83 |
| Goat anti-Rabbit IgG (H+L) Secondary Antibody, Alexa Fluor 488 | Invitrogen | #A11034 |
| Goat anti-Mouse IgG (H+L) Secondary Antibody, Alexa Fluor 568 | Invitrogen | #A |
| RNAscope Multiplex Fluorescent Detection Kit v2 | ACD | #323110 |
| RNAscope probe for mouse Slc48a1 | ACD | #585041-C2 |
| RNAscope probe for mouse SpiC | ACD | #1057881-C1 |
| RNAscope probe for mouse Hbb-bt | ACD | #540021-C3 |
| Annexin-V-Pacific Blue | BioLegend | #640918 |
| 7-AAD | eBioscience | #00-6993-50 |
| **Chemical and recombinant proteins** | **Source** | **Catalog number** |
| Hemin | Frontier Scientific | #H-651-9 |
| Sodium butyrate | Sigma-Aldrich | #B5887 |
| Human erythropoietin (EPO) | Amgen | #55513-144 |
| PolyJet in vitro DNA transfection reagent | SignaGen | #SL100688 |
| RPMI | Gibco | #11875-085 |
| TRIzol | Invitrogen | #15596018 |
| **Commercial assays** | **Source** | **Catalog number** |
| RNAscope multiplex fluorescent V2 assay kit | Advance Cell Diagnostics | Cat#323100 |
| SuperSignal West Pico PLUS chemiluminescent substrate | Thermo Scientific | Cat#3580 |
| SuperSignal West Femto maximum sensitivity substrate | Thermo Scientific | Cat#34095 |
| ProLong Diamond antifade mountant with DAPI | Invitrogen | Cat#P36962 |
| 10x Genomics Chromium X NextGEM 3' V3.1 Gel bead kit | 10x Genomics | N/A |
| KAPA Library Quantification Kit (Complete, Universal) | Kapa Biosystems | N/A |
| **Cell lines** | **Source** | **Identifier** |
| Human: K562 | ATCC | CCL-243 |
| **Mouse model** | **Source** | |
| WT: C57BL/6N-*Hrg1*$^{+/+}$ (HRG1-littermates) | This paper | N/A |
| *HRG1*-KO: C57BL/6N-*Hrg1*$^{em1(M10)Cas9}$ | This paper | N/A |
| *HRG1*-GFP: C57BL/6N-*Hrg1*$^{em1(TurboGFP)Cas9}$ | This paper | N/A |

| | | |
|---|---|---|
| Hbb$^{th3/+}$: B6;129-*Hbb-b1$^{tm1Unc}$ Hbb-b2$^{tm1Unc}$*/J | Jackson lab | Strain #:003253 |
| Hbb$^{th3/+}$;HRG1-KO: B6;129-*Hbb-b1$^{tm1Unc}$ Hbb-b2$^{tm1Unc}$*/J x C57BL/6N-*Hrg1$^{em1(M10)Cas9}$* | This paper | N/A |
| Hbb$^{th3/+}$;HRG1-GFP: B6;129-*Hbb-b1$^{tm1Unc}$ Hbb-b2$^{tm1Unc}$*/J x C57BL/6N-*Hrg1$^{em1(M10)Cas9}$* | This paper | N/A |
| **Oligonucleotides** | **Source** | **Identifier** |
| guide RNA for mouse strain HRG1$^{-/-}$: 5'-TAGGGACGGTGGTCTACCGACAACCGG-3' | This paper | N/A |
| guide RNA for mouse strain HRG1$^{tgfp/tgfp}$: 5'-CCTTAGTGATTTCTAACCCAGGG-3' | This paper | N/A |
| **Software and algorithms** | **Source** | **Identifier** |
| ImageJ | NIH/LOCI | https://imagej.net/ij/index.html |
| FlowJo version 10 | Tree Star | https://www.flowjo.com/ |
| Illumina's bcl2fastq2 Conversion Software v2.20 | Illumina | https://support.illumina.com/downloads/ bcl2fastq-conversion-software-v2-20.html |
| Imaris | Oxford Instruments | https://imaris.oxinst.com/ |
| R | R core team | https://www.R-project.org |
| RStudio | Posit team | https://posit.co/download/rstudio-desktop/ |
| Package: DESeq2 | Love et al.(*2*) | https://bioconductor.org/packages/release/bioc/html/DESeq2.html |
| Package: Seurat v5.2.1 | Hao et al (*3*), Hao et al.(*4*), Stuart et al.(*5*), Butler et al.(*6*), Satija et al.(*7*) | https://cran.r-project.org/web/packages/Seurat/ |
| Package: DoubletFinder v2.0.4 | McGinnis (*8*) | https://github.com/blaserlab/doubletfinder |
| Package: SingleR v2.6.0 | Aran et al. (*9*) | https://bioc.r-universe.dev/SingleR |
| **Other** | **Source** | **Identifier** |
| Hidex AMG automatic gamma counter | LabLogic Systems | N/A |
| IDEXX system (Westbrook, Maine, U.S.A.) | IDEXX Laboratories, Inc. | N/A |
| Synergy Neo2 plate reader | Agilent | N/A |
| Nikon W1 spinning disk confocal microscope | Nikon | N/A |
| Azure imaging systems | Azure | N/A |
| Canto | BD Biosciences | N/A |
| LSRII | BD Bioscience | N/A |
| Aurora3 | Cytek | N/A |
| AriaII | BD Biosciences | N/A |
| Countess 3 | Invitrogen | N/A |
| GX Touch (Revvity, Waltham, MA) | Revvity | N/A |
| Illumina 622 NovaSeq 6000 | Illumina | N/A |

### Methods
#### Animal studies

All mice were housed in a constant 12-hour light-dark cycle in the animal facilities at the University of Baltimore School of Medicine. They were given free access to tap water and a standard laboratory mouse diet. Males and females between 8 and 12 weeks old were used for experiments, but were plotted in separate graphs. *HRG1*-knockout ($Hrg1^{-/-}$, M10) mice were established on a C57BL/6N background through CRISPR/Cas9-mediated editing of exon 1, as reported previously (21). The M10 allele contains a two–base pair insertion and a single–base pair deletion, producing a frameshift mutation after the 37th amino acid in the Hrg1 coding sequence.(*1*) *HRG1-KO* mice were genotyped from tail genomic DNA extracts using a custom-ordered TaqMan SNP Genotyping Assays probe (ThermoFisher Scientific) on a Bio-Rad CFX Duet system.

HRG1-GFP mice were genotyped from the tail at P21. They were genotyped by PCR using primers HRG1-*Turbo*GFP Forward *5'*-TACTTAGCCTCTACGCCC-*3'*, *HRG1-TurboGFP* reverse 5'-TGCTCTTCATCTTGTTGGTCATGCGGC-*3',* and HRG1-*Turbo*GFP reverse 5'-TAACGGGCAGATTGGGCAGATTGGC-*3'*. β-thalassemic (B6.129P2-Hbb-$b1^{tm1Unc}$ Hbb-$b2^{tm1Un}$c/J /J, named as $Hbb^{Th3/+}$) mice were purchased from the Jackson Laboratory. $Hbb^{th3/+}$ mice were genotyped by qPCR using primers $Hbb^{th3/+}$ Forward *5'*-TACTGCCTGACCAAGGAAAGC-*3'*, $Hbb^{Th3/+}$ reverse *5'*-TGCTGACCTGCTGGATTACAT-*3'*. $Hbb^{th3/+}$ males were crossed to WT, HRG1-KO, HRG1-GFP females, and inbred to obtain the genotypes needed. WT mice denote wild-type littermates of each mouse strain. The Institutional Animal Care and Use Committee at the University of Maryland Baltimore approved all subsequent protocols involving these animals (IACUC Animal Study Protocol 1022007).

**Generation of KI (knock-in) mice**

Guide and Cas9 RNAs: The guide RNA (*5'- CCTTAGTGATTTCTAACCCAGGG -3'*) was purchased from Sage Laboratories 2033 Westport Center Drive, St Louis, MO. Cas 9 RNA was purchased from Trilink Biotechnologies, San Diego, CA. The repair template was generated by synthesizing the TurboGFP sequence with 250 bp homologous arms before and after the stop codon of mouse HRG1 in the pUC57 vector (Genscript), purified by the 2x CsCl method (Lofstrand Labs, Ltd), and linearized with NheI + XbaI enzymes. The guide RNA and Cas9 RNA were combined at a concentration of 5 ng/µl (each) in 10 mM Tris, 0.25 mM EDTA (pH 7.5) for injection. Pronuclear Injection: Pronuclear injection was performed using standard procedures(*10*). Briefly, fertilized eggs were collected from super ovulated C57BL/6J females approximately 9 h after mating with C57BL/6J male mice. Pronuclei were injected with a capillary needle with a 1–2 µm opening pulled with a Sutter P-1000 micropipette puller. The RNAs were injected using a FemtoJet 4i (Eppendorf) with continuous flow estimated to deposit approximately 2 pl of solution. Injected eggs were surgically transferred to pseudo-pregnant CB6 $F_1$ (BALB/cJ x C57BL/6J) recipient females. Genotyping: DNA was obtained from founder ($F_0$) animals by tail biopsy, amplified by PCR (Forward *5'-CCATCACAGTGTCTTTCTGAAGGG-3'* Reverse *5'-AGACCACACTCGTCCTGCTG -3'*), and sequenced to determine the insertion of TurboGFP. $F_0$ animals carrying TurboGFP insertion were crossed to C57BL/6J animals, and the resulting heterozygous F1 animals were either intercrossed (F) to generate homozygous KI animals or backcrossed (N) to C57BL/6J mice for propagation.

**Stress erythropoiesis models: phlebotomy (Px) and erythropoietin (EPO)**

Mice were bled 300 µL on two successive days (days 1 and 2) using the submandibular blood collection technique and subsequently intraperitoneally injected with 300 µL of saline to maintain the blood volume. EPO (200 UI, Amgen, #55513-144) was administered by intraperitoneal injection twice (days 1 and 2). Animals were euthanized 24 hours after the second phlebotomy or the second injection of EPO using a lethal dose of ketamine, then followed by cardiac perfusion using Dulbecco's phosphate-buffered saline (DPBS) (Gibco, cat. number 14190250). Prior to perfusion, whole blood was collected.

**Hematological parameters**

Blood was collected after sacrifice from the abdominal aorta in EDTA-coated tubes. Complete blood counts (CBCs) were assessed on an IDEXX system (Westbrook, Maine, U.S.A.).

**Flow cytometry**

Erythroblasts were analyzed from the bone marrow and the spleen by flow cytometry. Cell suspensions were washed and incubated on ice for 10 minutes in PBS-5% FBS with 0.1 µg of anti-mouse antibodies as follows: anti-Ter119-APCeFluor780 (Invitrogen, 47-5921-82), anti-CD44-PECy7 (Biolegend, 103030) or CD44-eFluor450 (Invitrogen, 40-0441-82), and for the lineage (Lin) PE-conjugated anti-Gr-1 (Invitrogen, 12-5931-83), anti-Cd4 (Invitrogen, 12-0041-82), anti-CD8a (Invitrogen, 12-0081-83), anti-CD45R (Invitrogen, 12-0452-83). Cells were acquired using Canto (BD Biosciences), LSRII (BD Bioscience), and Cytek Aurora3 (Cytek). Results were analyzed using Flow-Jo software (Tree Star, Ashland, OR).

For fluorescence-activated cell sorting (FACS) analysis, bone marrow from both the tibias and femurs and half of the spleen were washed and incubated on ice for 15 minutes with 0.1µg of anti-mouse antibodies: anti-Ter119-APCeFluor780 (Invitrogen, 47-5921-82), anti-CD44-eFluor450 (Invitrogen, 40-0441-82), and for the lineage (Lin) PE-conjugated anti-Gr-1-PE (Invitrogen, 12-5931-83), anti-Cd4-PE (Invitrogen, 12-0041-82), anti-CD45R (Invitrogen, 12-0452-83), anti-CD8a-PE (Invitrogen, 12-0081-83). Either Ter119-expressing cells or populations II to V were sorted for downstream analysis. Cells were sorted using Aria II (BD Bioscience). Apoptosis in erythroblasts was determined by flow cytometry using Annexin-V-Pacific Blue (BioLegend, 640918) and 7-AAD (eBioscience, 00-6993-50). The gating strategy is shown in Supplemental Figures S1A and S2D.

**Immunoblots**

Sorted erythroblasts were centrifuged at 300g for 5 minutes at 4°C and lysed with 15µL of lysis buffer (20 mM Hepes, 150 mM NaCl, 0.5% Triton-X100 + 1X protease inhibitor). Mammalian cells were lysed using a standard lysis protocol(*11*). The total protein concentration was measured using Bradford's assay, as described previously(*12*). The number of cells loaded is indicated on the immunoblots and used for normalization, as erythroblasts lose their organelles and become filled with hemoglobin as they mature; the protein content differs significantly across differentiated stages and cannot be used as a normalization method. Membranes were stripped before probing to remove heme and then probed overnight at 4°C in a blocking buffer containing the primary antibody of interest: mouse anti-HRG1(*1, 13*) (1:100), anti-Ter119 (eBioscience, #14-5921-85, 1:1000), or anti-tubulin (DSHB, #AA4.3, 1:1000). Following three rinses in PBST (PBS-tween 0.05%), membranes were exposed to horseradish peroxidase–linked secondary antibodies diluted 1:10,000 in 5% milk/PBST for one hour at ambient temperature. The membranes were subsequently washed three additional times with PBST and visualized using either Pico or Femto enhanced chemiluminescence substrates (Pierce) using Azure imaging systems. Quantification of the blots was performed using ImageJ and normalized by the number of cells counted or sorted, with Ter119 serving as a control for erythroblasts.

**Single-cell RNA-Seq**

100,000 cells of Ter119-containing cells (as previously described above in "Flow cytometry") were sorted from a WT bone marrow mouse. Cells were washed in 1X DPBS and pelleted. Cells were counted and viability assessed using a Countess 3 (Invitrogen, Carlsbad,CA). Cell suspensions were loaded onto a 10x Genomics Chromium X (Pleasanton,CA) using the NextGEM 3' V3.1 Gel bead kit. Cell capture, GEM generation, cDNA amplification, and library preparation were performed according to manufacturer protocol. The resulting 3' gene expression libraries were assessed for concentration and fragment size using the DNA High Sensitivity Assay on a GX Touch (Revvity, Waltham, MA). The libraries were pooled, assessed by qPCR using the KAPA Library Quantification Kit (Complete, Universal) (Kapa Biosystems, Woburn, MA), and sequenced on an Illumina NovaSeq 6000 using 100bp PE reads (Illumina, San Diego, CA). The resulting sequence data were demultiplexed using Illumina's bcl2fastq2 Conversion Software v2.20. Each sample was analyzed using 10x Genomics Cell Ranger v8.0.0(*14*). The reference for gene expression was refdata-gex-GRCh38-2024-A, available at https://www.10xgenomics.com/support/software/cell-ranger/downloads.

Raw sequencing data were processed using the Cell Ranger pipeline to generate gene-barcode matrices. Initial quality control was performed in Seurat (v5.2.1). Cells with fewer than 3 detected genes and genes expressed in fewer than 5 cells were excluded. Additional mitochondrial filtering criteria included total RNA counts greater than 5, fewer than 2,500 detected features per cell (nFeature_RNA < 2500), and mitochondrial gene content below 5%. Doublets were removed using the DoubletFinder package (v2.0.4). The parameters used were pN = 0.05, pK = 0.3 for WT, with the optimal pK determined using the find.pK() function applied to sweep statistics generated by paramSweep() and summarizeSweep(). The expected number of doublets (nExp) was estimated as 2.3% of the total number of cells. Datasets were then integrated using Seurat to correct for potential batch effects. Marker genes and clusters were identified using the FindAllMarkers() function with the following parameters: logfc.threshold = 0.25, min.pct = 0.5, only.pos = TRUE, test.use = 'DESeq2', and slot = 'counts'. Cell type annotation was performed using the SingleR package (v2.6.0).

**RNA in situ hybridization using RNAscope**

Femora were fixed in 2% formaldehyde solution for 2 days and then embedded in OCT compound (Fisher Scientific). Frozen sections of 30µm thickness were made on the Kawamoto film (SECTION-LAB)(*15*). RNAscope was performed on the Kawamoto film using the RNAscope Multiplex Fluorescent V2 Assay Kit (Advance Cell Diagnostics) as previously described(*16, 17*). Probes for *Hbb-bt* (ACD, #540021-C3), *Spic* (ACD, #1057881-C1), and *Slc48a1* (ACD #585041-C2) were visualized using OPAL520, OPAL570, and OPAL690, respectively (Akoya Biosciences). Images were captured with a CSU-W1 spinning disk confocal microscope (Nikon) at 0.2 µm steps using a 60x objective with oil. Images were reconstructed using Imaris (Oxford Instruments).

**Mammalian cell culture, transfection, treatments**

Mammalian cell culture experiments were performed using K562 cells WT and *HRG1*-KO. These cells were grown in a humidified incubator at 37°C with 5 % CO2. RPMI1640 (Gibco, #11875-

085) supplemented with 10% HI-FBS (heat-inactivated Fetal Bovine Serum, GeminiBio, #100-106 A424041), 1% Penicillin-Streptamycin-Glutamine (PSG, Gibco, #10-378-016), and 1% non-essential amino acids (NEAA, Gibco, #11140-050) was used as the culture medium. Transfections for immunoblotting, reporter assays, immunofluorescence, and imaging were done using PolyJet (SignaGen, SL100688).
Hemin chloride (Frontier CAS# 16009-13-5) was dissolved in a 1:1 (v/v) mixture of ethyl alcohol and 0.2 M KOH to obtain a 10mM final concentration.

Sodium butyrate (Sigma-Aldrich, #B5887) was dissolved in DPBS 1X.

**Generation of *HRG1*-KO cells**
SgRNA guide sequences targeting human HRG1 were cloned into the pSpCas9-2A-GFP plasmid as described by Ran et al(*18*). One million K562 cells were grown in RPMI supplemented with 10% FBS, 1% PSG, and 1% NEAA and plated in a 6-well plate. Cells were transfected with pSpCas9-2A-GFP carrying sgRNAs using PolyJet (Signagen, SL100688). GFP-positive cells were sorted by flow cytometry in a 96-well plate and validated by sequencing using forward (*5'-TTCGGGCCCTGCACCTGTGACTCTCG -3'*) and reverse (*5'- TCTGTGCAGCCGGCAGTGG -3'* for exon 1, and *5'- AGGTTTCCTCGAAAGGGGACG -3'* for exon 1 to 3) primers. K562 cells were harvested and washed with 1X DPBS once and then resuspended to a final volume of 100 µL. The pellet was sonicated with a Bioruptor on the high setting, with 30 seconds on and 30 seconds off for two 15-minutes cycles. Next, 50 µL of Proteinase K (10 mg/mL) was added, followed by incubation at 65°C for one hour, until the solution was clear. Next, 20 µL of RNaseA (10 mg/mL) was added, and then the solution was incubated at 37°C for 30 minutes. Following incubation, the DNA was purified using the Qiagen PCR purification kit and eluted with 30 µL of TE (Tris-EDTA) buffer. Quantification of DNA was done with PicoGreen (ThermoFisher).

***O*-dianisidine staining**
*O*-dianisidine staining was performed to detect hemoglobin in maturing K562 WT and *HRG1*-KO cells, as previously described(*13, 19*). Briefly, K562 cells were placed in 200µL of staining solution (0.6 (w/v) *O*-dianisidine, 25% ethanol, 10 mM sodium acetate, and 0.02% $H_2O_2$) for 10 minutes in dark conditions. Heme catalyzes the oxidation of *o*-dianisidine in the presence of $H_2O_2$, producing a dark brown color in hemoglobin-positive cells. Total cells and positive cells were counted using a hemocytometer.

**RNA-Seq**
K562 WT and *HRG1*-KO cells were grown on T75 flasks with RPMI supplemented with 10% HI-FBS, 1% PSG, and 1% non-essential amino acids. Cells were not treated (control) or treated with 0.75 mM sodium butyrate (SB) 24 hours after being plated and treated a second time 48 hours after the first treatment. Cells were collected 6h after the second treatment and washed three times with 1X DPBS. Total RNA was extracted using TRIzol lysis reagent (Invitrogen, #15596018). A total of 12 samples were sequenced using Illumina's Novaseq platform (Novogene), with a triplicate of each genotype and condition. Paired-end 150 base reads were sequenced and checked by FastQC, version 0.12.1. The reads were aligned to the human reference genome hg38 (UCSC) using STAR alignment (v2.7.11b). Differentially expressed genes (DEGs) were identified using DESeq2 (v1.44.0) in R (v4.4.1). Log2 fold changes in gene expression were shrunk using the lfcShrink function with the 'ashr' method. Adjusted p-values were calculated using the Benjamini-Hochberg (BH) method, with a significant cutoff of 0.05. The volcano plot was generated using the

EnhancedVolcano package (v0.99.5). Enrichment analysis was performed with significantly differentially expressed genes ($p < 0.05$, $Log_2FC > 1$) using Enrichr (Ma'ayan Laboratory).

**Immunofluorescence of mammalian cells**

K562 cells were fixed using 4% paraformaldehyde (PFA), blocked in 5% BSA (bovine serum albumin), and incubated with primary antibodies anti-HRG1 (1:100) and anti-CD71 (BioPharmingen, #55134), followed by secondary antibodies anti-rabbit 488 (1:5000) and anti-mouse 568 (1:5000). Cells were then washed with PBS thrice and subsequently mounted using Antifade (ProLong Gold Antifade Reagent, Invitrogen, CAS P36934). Images were taken and processed using Nikon A1 confocal microscope. Images have been denoised using Nikon NIS-elements software.

**Live cell imaging**

K562 cells were transfected with hHRG1-GFP using PolyJet (SignaGen, SL100688). 16 hours post-transfection, cells were transferred to Ibidi-coated dishes (check cat no) and allowed to lay down. Cells were then imaged using a Nikon W1 spinning disk microscope after 16 hours. Videos were processed and denoised by Nikon NIS-Elements software.

**Image analysis and 3D rendering**

All image analysis was performed using Imaris software (Bitplane). 3D rendering was also done using Imaris (Bitplane).

**[$^{59}$Fe]-heme uptake**

[$^{59}$Fe]-heme was synthesized by incorporating 18mCi/mg $^{59}FeCl_3$ (Perkin Elmer) into protoporphyrin IX (Frontier Scientific) as previously outlined(*20*). K562 WT or KO cells were seeded at 25,000 cells/ml in a 6-well plate in duplicate or triplicate. Cells were not treated (control) or treated with 0.75 mM sodium butyrate 24 hours after being plated. 48 hours after the first treatment, cells were treated again 48 hours later with sodium butyrate and with radiolabeled $^{59}$Fe-Heme and incubated either at 4°C for 6 hours or 37°C for 24 hours. The ratio between 37°C and 4°C reflects heme accumulation within cells while removing the heme-binding effect.

**Bioinformatics and statistics**

All data are presented as mean ± the standard error of the mean (SEM). Statistical significance was determined using one-way or two-way ANOVA or unpaired t test, as indicated in figure legends using GraphPad Prism, version 10.00 (GraphPad Software, Inc). Outliers were removed using "Outlier Calculator" at https://MiniWebtool.com//outlier-calculator// using the Moore and McCabe method(*21*).

**Fig. S1.**

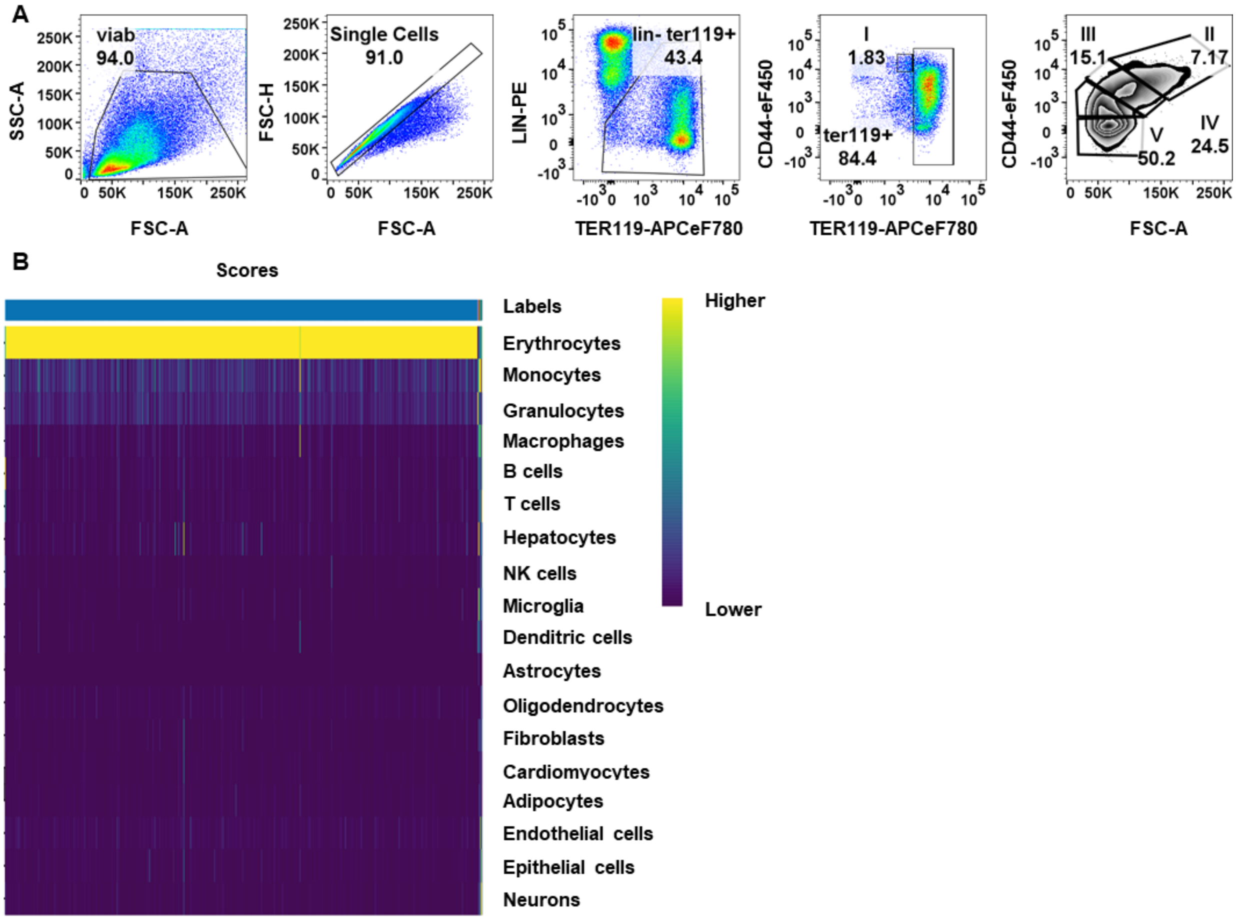


**Figure S1. Flow cytometry gating strategy and lineage annotation of single-cell RNA-seq data from WT bone marrow erythroblasts.**(**A**) Representative gating strategy to assess the maturation of erythroblasts. Data from Figure 2D is used as an example. Live cells were first identified based on forward scatter area (FSC-A, cell size) and side scatter area (SSC-A, cell granularity), followed by singlet discrimination using FSC-H (forward scatter height) versus FSC-A. Lineage-negative cells (excluding immune cells) were gated and Ter119$^{+}$ cells were selected to enrich for erythroblasts. Erythroid populations were then resolved based on Ter119 and CD44 expression, allowing discrimination of population I (proerythroblasts; CD44$^{high}$Ter119$^{medium}$) from populations II-V (Ter119$^{high}$). Forward scatter was subsequently used to distinguish the four stages of erythroid maturation: population II, basophilic erythroblasts; population III, polychromatic erythroblasts; population IV, orthochromatic erythroblasts and reticulocytes; and population V, mature red blood cells. (**B**) Heatmap showing cell-type annotation scores for all cells captured by single-cell RNA sequencing following erythroblast sorting from the bone marrow of WT mice. Each column represents a single cell and each row corresponds to reference cell lineages. Vertical bar depicts relative confidence score for each lineage assignment with the highest scores colored in yellow.

Fig. S2.

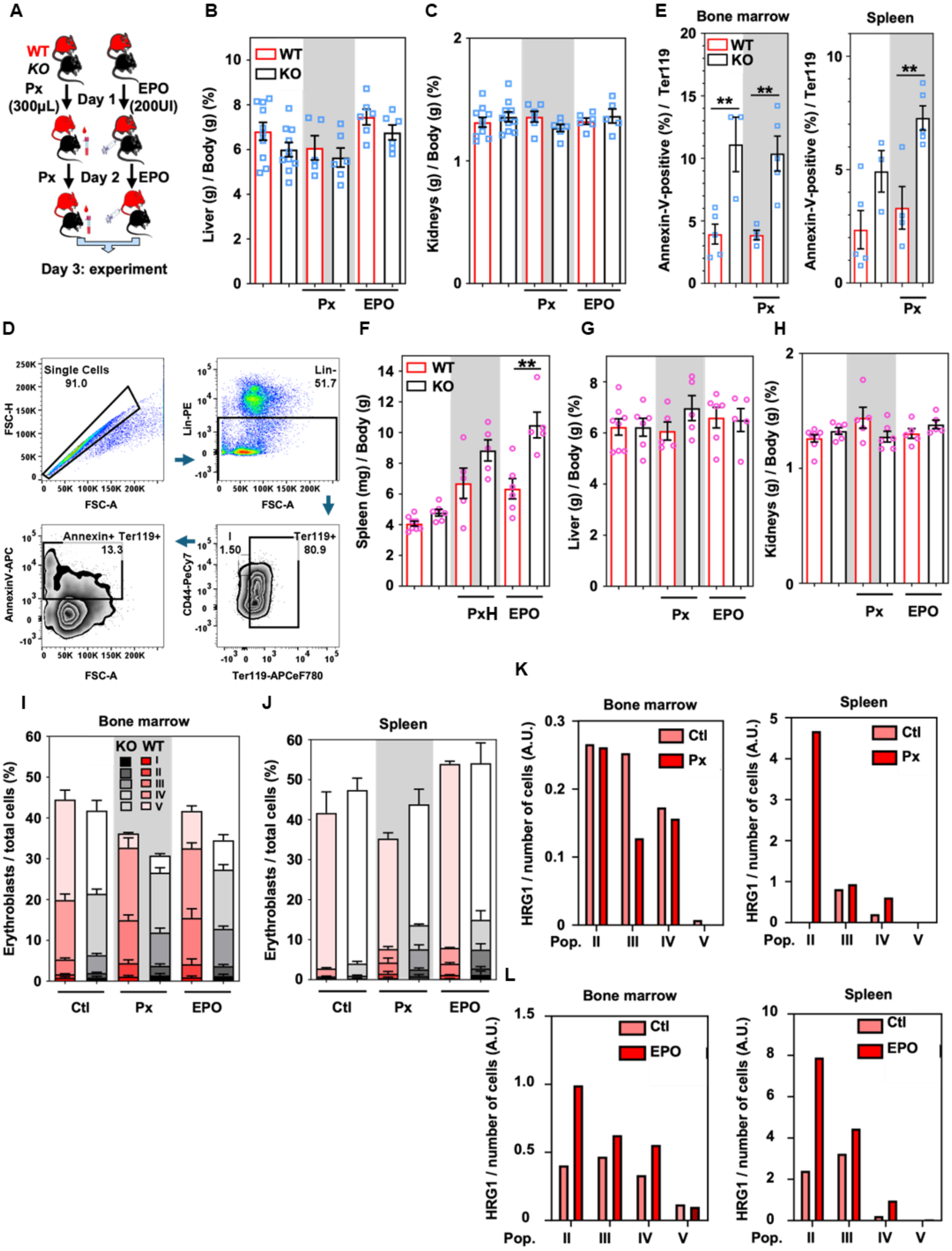


**Figure S2. HRG1 deletion induces the same trend of erythropoiesis defect in female mice.**

(**A**) Schematic of the experimental design of stress erythropoiesis models. WT (in red) or *HRG1-KO* (KO, in black) male mice were either phlebotomized (Px, 300µL) or injected with 300µL of saline, or injected with 200 U.I. of erythropoietin (EPO) on day 1 and day 2, 24 hours apart, and the experiment was performed 24 hours following the second treatment. The ratio of liver (**B**) and kidney (**C**) weight normalized to the body weight of WT and *HRG1-KO* male mice (open blue square) in control, phlebotomies, and EPO conditions. Error bars represent mean ± SEM, n=5-12 mice/group. (**D**) Representative gating strategy for Annexin-V positive cells. (**E**) The percentage of apoptotic Annexin-V positive cells in the Ter119-expressing erythroblasts in the bone marrow and spleen. Error bars represent mean ± SEM, n=3-5mice/group.*p<0.05, **p<0.01, ***p<0.001 (One-way ANOVA, Sidak's multiple comparison test). The ratio of (**F**) spleen ,(**G**) liver , and (**H**) kidney weight normalized to the body weight of WT and *HRG1-KO* female mice (open pink circle) in control, phlebotomies, and EPO conditions. Error bars represent mean ± SEM, n=5-12 mice/group. **p<0.01 (One-way ANOVA, Sidak's multiple comparison test). Flow cytometry analysis of the proportion of erythroblasts in the live single cell population in the (**I**) bone marrow and (**J**) the spleen of control, phlebotomized, or EPO-injected WT and *HRG1-KO* female mice (open pink circle). WT and HRG1-KO female mice from the same condition for each erythroid population were compared using an unpaired t-test, with no significant difference observed. The immunoblots were quantified from (**K**) Figure 2G and (**L**) Figure 2H and normalized to the number of cells loaded.

**Fig. S3.**

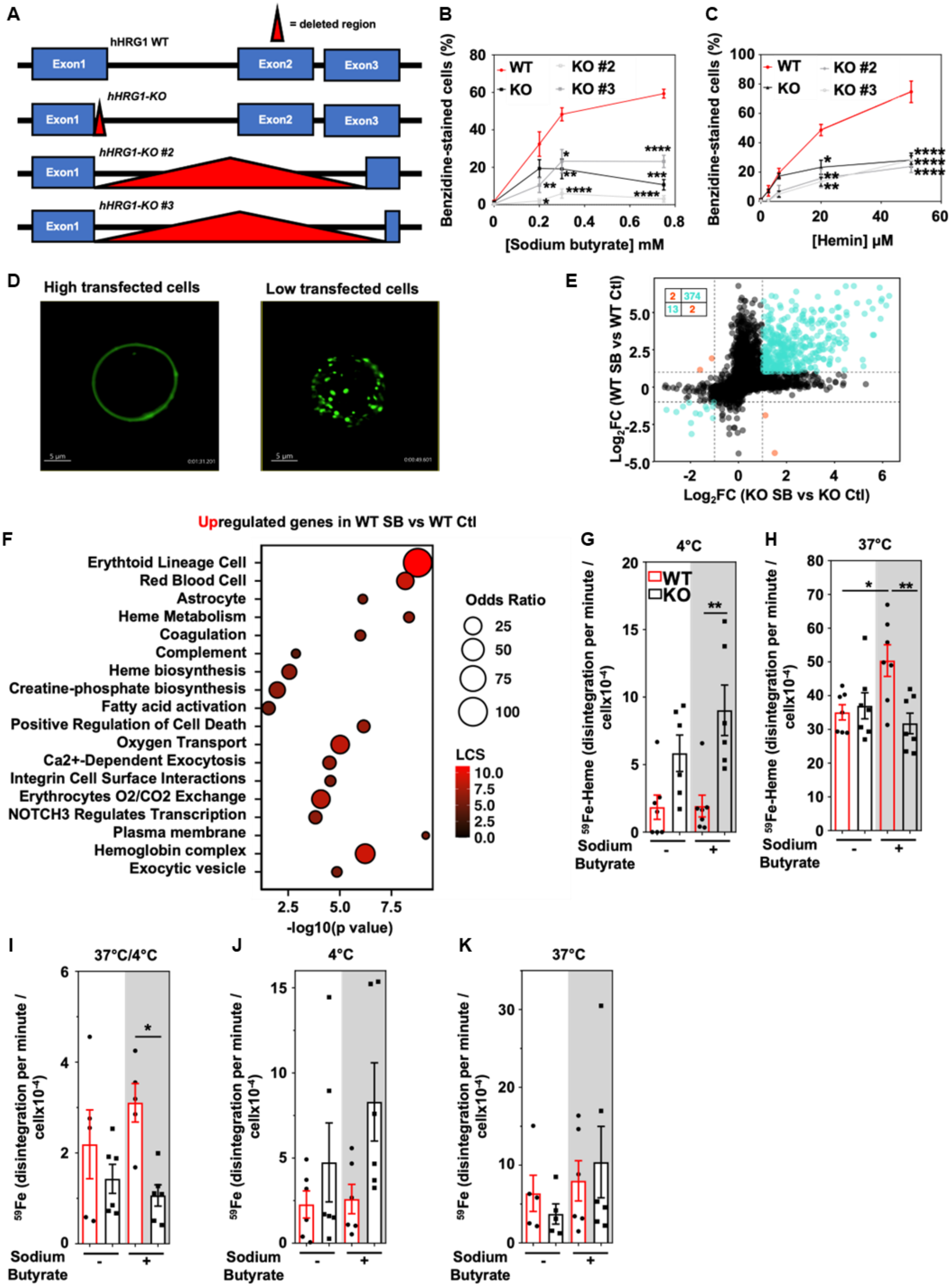
A
= deleted region
hHRG1 WT
Exon1
Exon2
Exon3
hHRG1-KO
hHRG1-KO #2
hHRG1-KO #3
B
Benzidine-stained cells (%)
WT
KO
KO #2
KO #3
[Sodium butyrate] mM
C
[Hemin] μM
D
High transfected cells
Low transfected cells
E
Log2FC (WT SB vs WT Ctl)
Log2FC (KO SB vs KO Ctl)
F
Upregulated genes in WT SB vs WT Ctl
Erythtoid Lineage Cell
Red Blood Cell
Astrocyte
Heme Metabolism
Coagulation
Complement
Heme biosynthesis
Creatine-phosphate biosynthesis
Fatty acid activation
Positive Regulation of Cell Death
Oxygen Transport
Ca2+-Dependent Exocytosis
Integrin Cell Surface Interactions
Erythrocytes O2/CO2 Exchange
NOTCH3 Regulates Transcription
Plasma membrane
Hemoglobin complex
Exocytic vesicle
-log10(p value)
Odds Ratio
LCS
G
4°C
59Fe-Heme (disintegration per minute / cellx10-4)
Sodium Butyrate
H
37°C
I
37°C/4°C
59Fe (disintegration per minute / cellx10-4)
J
4°C
K
37°C

**Figure S3. RNAseq confirms erythroid differentiation of K562 cells with sodium butyrate treatment.**

**(A)** Schematic of the deleted sequence in the HRG1 gene in K562 cells. In HRG1-KO mutants, 161 base pairs were deleted at the end of exon 1. In HRG1-KO #2 mutants, 6845 base pairs were deleted corresponding to the region between the end of exon 1 to the middle of exon 3. In hHRG1-KO #3 mutants, 6884 base pairs were deleted corresponding to the region between the end of exon 1 to the end of exon 3. Benzidine staining of WT and all mutants *HRG1*-KO K562 cells treated with different concentrations of (**B**) sodium butyrate (SB) or (**C**) hemin and harvested 72h after the second treatment. Error bars represent mean ± SEM from 3 biological replicates comprising at least 2 technical replicates. WT and *HRG1-KO* cells were analyzed for each time point using an unpaired t-test, *p<0.05, **p<0.01, ****p<0.0001. (**D**) Snapshot of live-cell imaging of K562 cells highly or lowly transfected with hHRG1-GFP. Scale bars as indicated. (**E**) Scatter plots of *HRG1-KO* SBvs. HRG1-KO control (ctl) (x-axis) against $Log_2$ fold chance ($Log_2FC$) of WT SB vs. WT control (ctl) (y-axis). Broken black lines mark $Log_2FC = 1$. (**F**) Enrichment analysis plots of the 678 genes that are upregulated in WT cells SB compared to WT control. K562 WT and *HRG1*-KO K562 cells were plated, treated with 0.75 mM sodium butyrate 24 hours later, treated again 48 hours later with sodium butyrate (SB) and with (**G-H**) radiolabeled $^{59}$Fe-Heme or (**I-K**) $^{59}$Fe and incubated either at (**G&J**) 4°C for 6h or (**H&K**) 37°C for 24 hours. (**I**) The ratio between 37°C and 4°C reflects iron accumulation within cells while removing the iron binding effect. Error bars represent mean ± SEM from 3 biological replicates comprising at least 2 technical replicates.*p<0.05, ***p<0.001 (One-way ANOVA, Sidak's multiple comparison test).

**Fig. S4.**

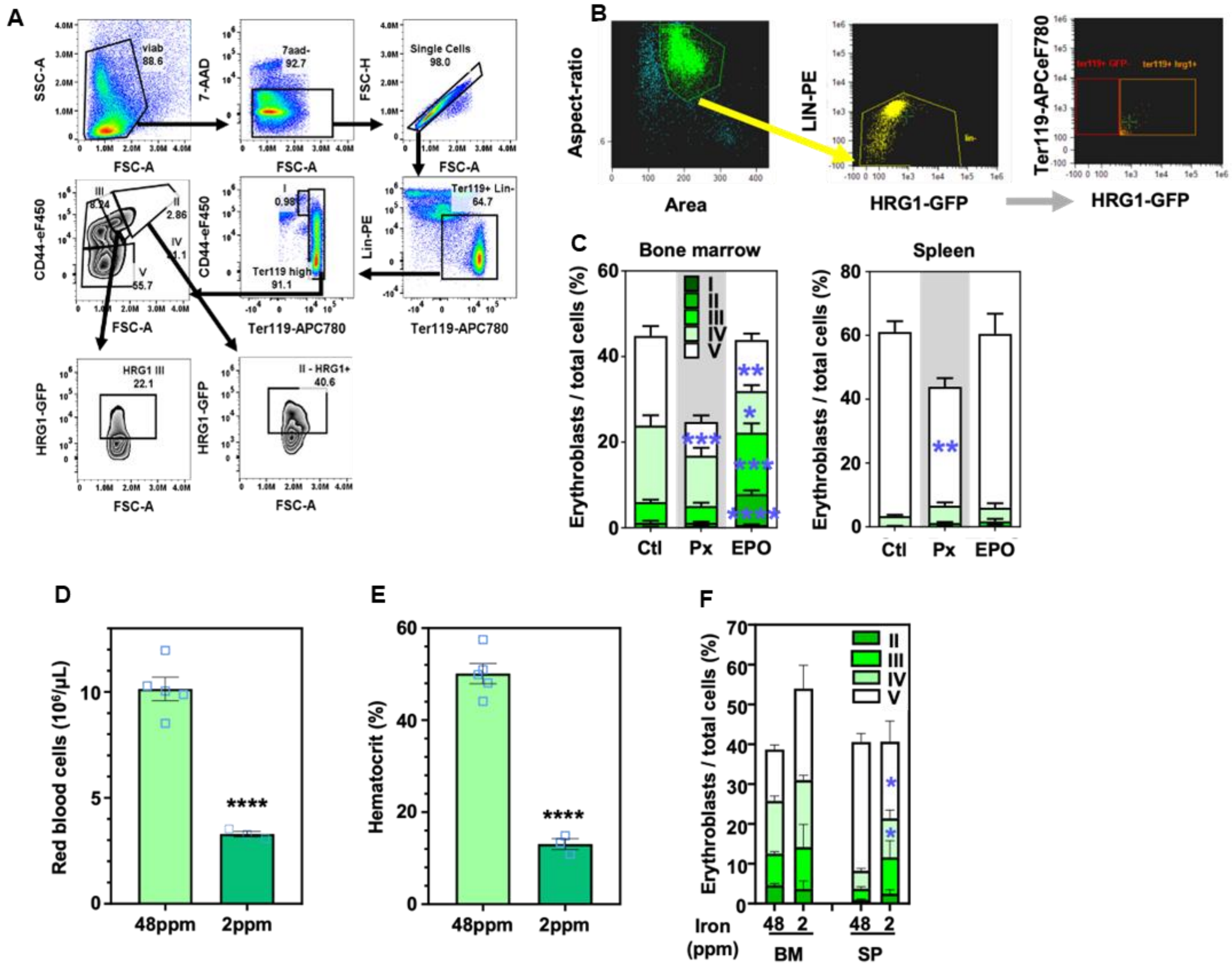


**Figure S4. Iron-deficiency diet induces anemia.**

(**A**) Representative gating strategy for HRG1-GFP-containing cells. (**B**) Representative gating strategy for HRG1-expressing cells using Flowsight. (**C**) Flow cytometry analysis of the proportion of erythroid precursors in the live single cell population in the bone marrow (left panel) and the spleen (right panel). Analysis was performed in control, phlebotomized (Px), or erythropoietin (EPO)-injected *HRG1*$^{GFP/+}$ mice. Error bars represent mean ± SEM, n=3-5 mice/group. Each erythroid population was compared between groups using an unpaired t-test between control and treated conditions. **p<0.01, ***p<0.001, ****p<0.0001. Blood parameters from *HRG1*$^{GFP/+}$ mice fed with an iron-deficient (IDD, 2 ppm) or -sufficient diet (ISD, 48 ppm) (**D**) red blood cell count and (**E**) hematocrit. Error bars represent mean ± SEM, n=5-10 mice/group. *<0.05, ***p<0.001, ****p<0.0001 (unpaired t-test). (**F**) Flow cytometry analysis of the proportion of erythroid precursors in the alive single cell population in the bone marrow (BM) and the spleen (SP), in *HRG1*$^{GFP/+}$ mice fed with a 2 ppm or 48 ppm iron diet. Error bars represent mean ± SEM, n=3-5 mice/group. Each erythroid population was compared between groups using an unpaired t-test between 48ppm and 2ppm conditions. *p<0.05.

**Fig. S5.**

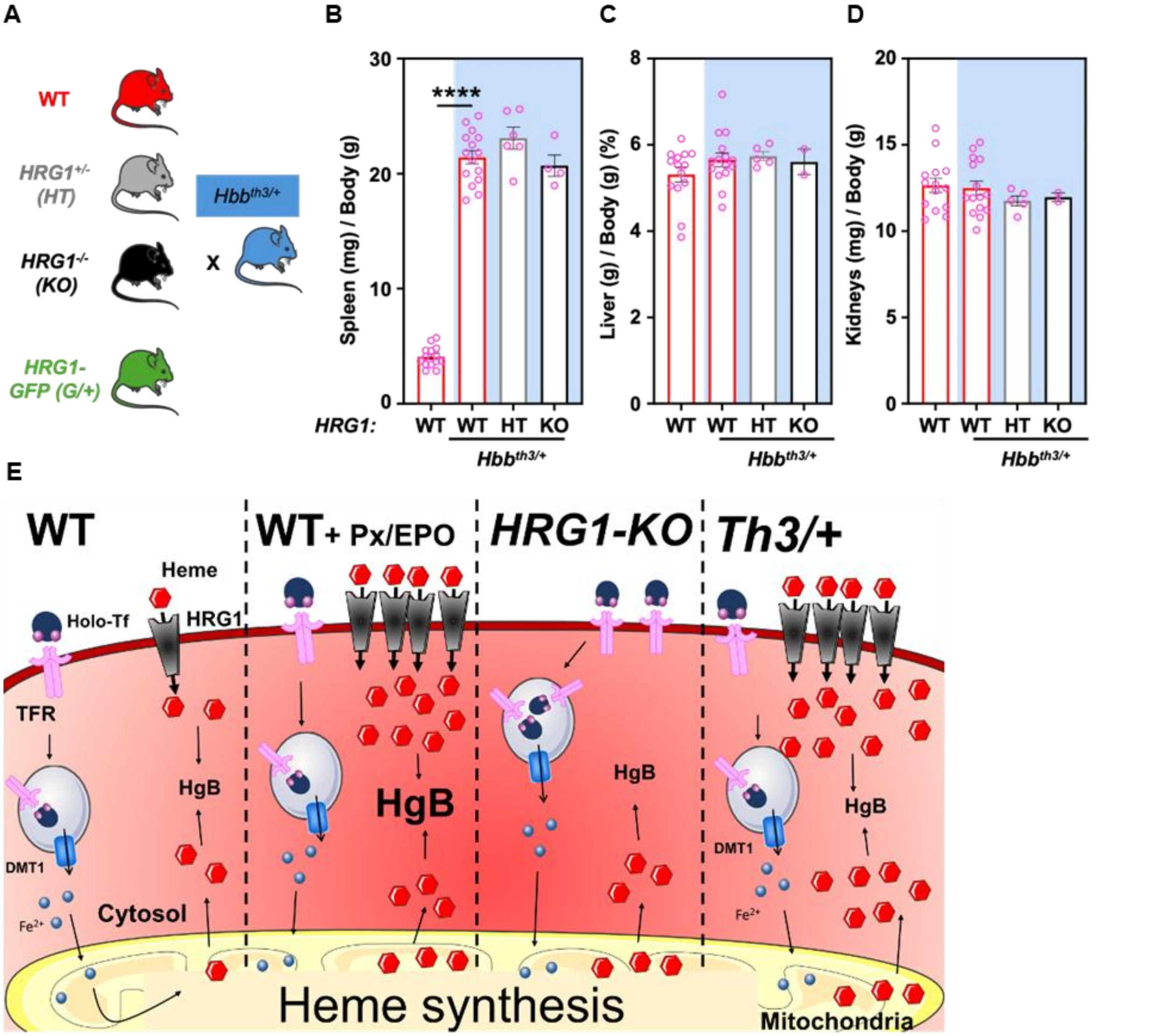


**Figure S5. HRG1 deletion did not exacerbate splenomegaly in β-thalassemic mice**

(**A**) Schematic of mice crosses generated: *Hbb*$^{th3/+}$ (β-thalassemic) mice were either crossed to WT, *HRG1-KO* (KO), *HRG1-GFP* (G/+). The ratio of (**B**) spleen, (**C**) liver, and (**D**) kidneys' weight normalized to the body weight of WT, *Hbb*$^{th3/+}$*, Hbb*$^{th3/+}$*;HRG1-HT* (one allele of HRG1 being removed)*,* and *Hbb*$^{th3/+}$*;HRG1-KO* female mice (open pink circle). Error bars represent mean ± SEM, n=3-12 mice/group. ****p<0.0001 (One-way ANOVA, Sidak's multiple comparison test). (**E**) Proposed model for exogenous heme import by HRG1 in erythroblasts at steady-state and during stress erythropoiesis in WT, *HRG1*-KO mice and β-thalassemia.

**Table S1.**

| Sex | Genotype | Treatment | RBC (M/µL) | HGB (g/dL) | HCT (%) | MCV (fL) | MCH (pg) | MCHC (g/dL) | RDW-SD (fL) | RDW-CV (%) | RETICS (K/µL) | RETICS (%) |
|---|---|---|---|---|---|---|---|---|---|---|---|---|
| Male | WT | Ctl | 10.13 ± 0.9 | 15.33 ± 1.43 | 49.21 ± 5.55 | 48.5 ± 1.99 | 15.12 ± 0.17 | 31.22 ± 1.17 | 28.22 ± 2.85 | 24.28 ± 1.7 | 533 ± 118 | 5.24 ± 0.89 |
| | | Px | 6.31 ± 0.81 | 9.66 ± 1.21 | 31.66 ± 3.08 | 50.36 ± 2.8 | 15.32 ± 0.15 | 30.48 ± 1.51 | 29.98 ± 5.22 | 23.3 ± 2.82 | 692 ± 115 | 10.93 ± 0.64 |
| | | EPO | 10.41 ± 0.5 | 15.68 ± 0.74 | 51.4 ± 3.06 | 49.33 ± 0.8 | 15.07 ± 0.12 | 30.55 ± 0.65 | 28.22 ± 1.31 | 27.05 ± 1.24 | 810 ± 100 | 7.78 ± 1.0 |
| | *HRG1-KO* | Ctl | 9.7 ± 1.07 | 14.56 ± 1.64 | 46.43 ± 5.43 | 47.86 ± 1.19 | 15.02 ± 0.26 | 31.37 ± 0.68 | 29.94 ± 2.67 | 25.35 ± 2.21 | 530 ± 85 | 5.48 ± 0.69 |
| | | Px | 4.72 ± 0.75 | 7.18 ± 1.17 | 23.35 ± 3.08 | 49.88 ± 6.24 | 15.18 ± 0.34 | 30.83 ± 3.34 | 27.28 ± 5.63 | 21.3 ± 2.08 | 546 ± 190 | 11.31 ± 2.67 |
| | | EPO | 11.05 ± 0.54 | 16.37 ± 0.79 | 52.13 ± 2.82 | 47.18 ± 1.14 | 14.8 ± 0.24 | 31.38 ± 0.95 | 29.78 ± 2.69 | 29.55 ± 1.32 | 763 ± 105 | 6.95 ± 1.26 |
| Female | WT | Ctl | 9.84 ± 1.25 | 15.14 ± 1.93 | 46.2 ± 6.14 | 47.02 ± 2.34 | 15.39 ± 0.27 | 32.86 ± 2.25 | 27.71 ± 3.06 | 24.56 ± 3.68 | 516 ± 98 | 5.3 ± 0.48 |
| | | Px | 5.6 ± 1.32 | 8.76 ± 2.06 | 27.98 ± 6.58 | 49.98 ± 0.64 | 15.64 ± 0.18 | 31.32 ± 0.63 | 27.7 ± 1.4 | 24.74 ± 2.37 | 873 ± 155 | 14.48 ± 2.61 |
| | | EPO | 10.78 ± 0.84 | 16.55 ± 1.46 | 53.52 ± 4.24 | 49.65 ± 0.8 | 15.35 ± 0.36 | 30.92 ± 0.82 | 29.77 ± 2.28 | 27.33 ± 1.89 | 874 ± 224 | 8.03 ± 1.73 |
| | *HRG1-KO* | Ctl | 10.31 ± 1.35 | 15.42 ± 2.31 | 49.1 ± 8.14 | 47.43 ± 1.78 | 14.93 ± 0.5 | 31.48 ± 0.95 | 29.75 ± 3.57 | 26.02 ± 2.35 | 557 ± 153 | 5.34 ± 0.98 |
| | | Px | 4.11 ± 1.37 | 6.44 ± 2.14 | 20.04 ± 5.34 | 49.74 ± 3.84 | 15.68 ± 0.31 | 31.7 ± 2.58 | 29.34 ± 6.26 | 26.2 ± 5.22 | 557 ± 225 | 13.09 ± 3.28 |
| | | EPO | 10.71 ± 1.02 | 16.38 ± 2.04 | 51.72 ± 5.25 | 48.38 ± 3.77 | 15.24 ± 0.66 | 31.76 ± 4.02 | 32.08 ± 6.0 | 30.46 ± 3.68 | 867 ± 225 | 8.23 ± 1.77 |

**Supplemental Table 1. Blood parameters from WT and HRG1-KO mice males and females in control, phlebotomies, and EPO-injected conditions.**
Values are represented as mean ± SEM from 5-12 mice/group.

**Table S2.**

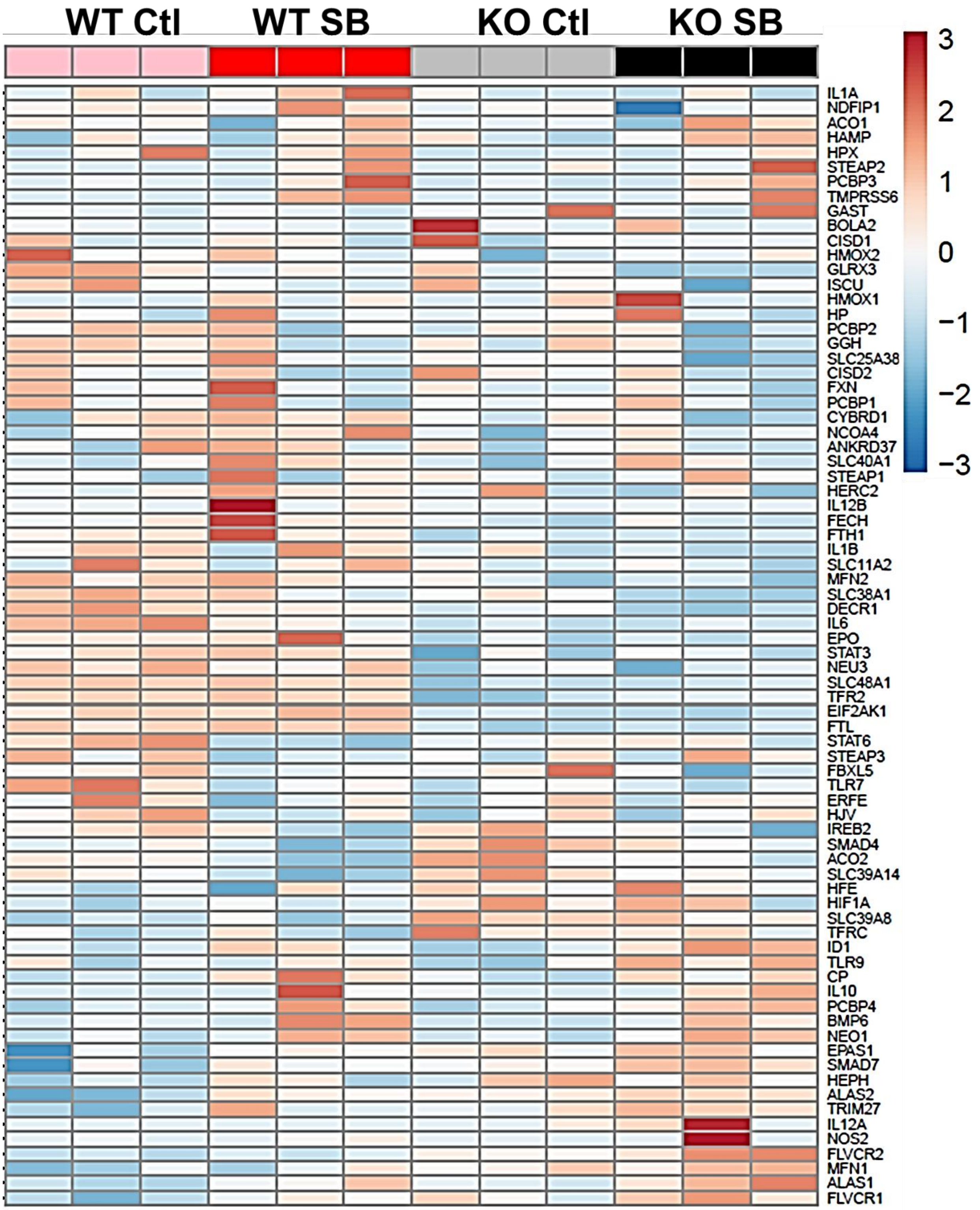


**Supplemental Table 2. Heme-iron genes heatmap.**
Heatmap of heme-iron genes in control and treated WT and HRG1-KO K562 cells with sodium butyrate.

#### Table S3

| Genotype Th3/+ | Genotype HRG1 | RBC (M/µL) | HGB (g/dL) | HCT (%) | MCV (fL) | MCH (pg) | MCHC (g/dL) | RETICS (K/µL) | PLT (K/µL) | WBC (K/µL) |
|---|---|---|---|---|---|---|---|---|---|---|
| WT | WT | 8.8 ± 1.81 | 13.02 ± 2.64 | 40.12 ± 8.69 | 45.48 ± 1.08 | 14.79 ± 0.21 | 32.55 ± 0.63 | 346 ± 102 | 450.77 ± 274.18 | 4.77 ± 2.87 |
| Th3/+ | WT | 5.61 ± 1.44 | 7.91 ± 2.15 | 24.83 ± 6.54 | 44.15 ± 2.83 | 14.09 ± 1.23 | 32.01 ± 2.92 | 1987 ± 479 | 910.08 ± 1193.39 | 13.92 ± 6.28 |
| | HT | 6.22 ± 1.05 | 8.72 ± 1.24 | 27.92 ± 4.51 | 44.97 ± 2.37 | 14.07 ± 0.78 | 31.35 ± 2.03 | 2358 ± 540 | 689.83 ± 154.23 | 21.34 ± 2.4 |
| | *KO* | 4.87 ± 1.37 | 7.17 ± 0.91 | 20.33 ± 5.18 | 42.03 ± 2.25 | 15.23 ± 3.02 | 36.07 ± 5.23 | 1576 ± 867 | 2496.0 ± 2778.22 | 16.54 ± 9.0 |

**Supplemental Table 3. Blood parameters from WT and *Hbb$^{th3/+}$;HRG1-WT*, *Hbb$^{th3/+}$;HRG1-HT, Hbb$^{th3/+}$;HRG1-KO* females.**
Values are represented as mean ± SEM from 3-15 mice/group.

#### Table S4

| BioProject | BioSample | SRA | Sample name |
|---|---|---|---|
| PRJNA1366593 | SAMN53317353 | SRR36103504 | C57Bl6_WT_BM |
| PRJNA1366593 | SAMN53347971 | SRR36127134 | WT control 1 |
| PRJNA1366593 | SAMN53347972 | SRR36127133 | WT control 2 |
| PRJNA1366593 | SAMN53347973 | SRR36127130 | WT control 3 |
| PRJNA1366593 | SAMN53347974 | SRR36127129 | HRG1-KO control 1 |
| PRJNA1366593 | SAMN53347975 | SRR36127128 | HRG1-KO control 2 |
| PRJNA1366593 | SAMN53347976 | SRR36127127 | HRG1-KO control 3 |
| PRJNA1366593 | SAMN53347977 | SRR36127126 | WT sodium butyrate 1 |
| PRJNA1366593 | SAMN53347978 | SRR36127125 | WT sodium butyrate 2 |
| PRJNA1366593 | SAMN53347979 | SRR36127124 | WT sodium butyrate 3 |
| PRJNA1366593 | SAMN53347980 | SRR36127123 | HRG1-KO sodium butyrate 1 |
| PRJNA1366593 | SAMN53347981 | SRR36127132 | HRG1-KO sodium butyrate 2 |
| PRJNA1366593 | SAMN53347982 | SRR36127131 | HRG1-KO sodium butyrate 3 |

**Supplemental Table 4. BioSample and SRA run accessions for each sample of the BioProject accession PRJNA1366593.**
